# Comprehensive Analysis of Spatial Architecture in Primary Liver Cancer

**DOI:** 10.1101/2021.05.24.445446

**Authors:** Rui Wu, Wenbo Guo, Xinyao Qiu, Shicheng Wang, Chengjun Sui, Qiuyu Lian, Jianmin Wu, Yiran Shan, Zhao Yang, Shuai Yang, Tong Wu, Kaiting Wang, Yanjing Zhu, Shan Wang, Changyi Liu, Yangqianwen Zhang, Bo Zheng, Zhixuan Li, Yani Zhang, Siyun Shen, Yan Zhao, Wenwen Wang, Jinxia Bao, Ji Hu, Xuan Wu, Xiaoqing Jiang, Hongyang Wang, Jin Gu, Lei Chen

**Author notes:** Equal Contribution (Rui Wu, Wenbo Guo, Xinyao Qiu). Corresponding authors: Lei Chen, Ph.D. International Cooperation Laboratory on Signal Transduction, Eastern Hepatobiliary Surgery Institute, 225 Changhai Road, Shanghai 200438, China. Jin Gu, Ph.D. MOE Key Laboratory for Bioinformatics, BNRIST Bioinformatics Division, Department of Automation, Tsinghua University, Beijing 100084, China. Hongyang Wang, M.D. International Cooperation Laboratory on Signal Transduction, Eastern Hepatobiliary Surgery Institute, 225 Changhai Road, Shanghai 200438, China.

## Abstract

Heterogeneity is the major challenge for cancer prevention and therapy. Here, we firstly constructed high-resolution spatial transcriptomes of primary liver cancers (PLCs) containing 84,823 spots within 21 tissues from 7 patients. The sequential comparison of spatial tumor microenvironment (TME) characteristics from non-tumor to leading-edge to tumor regions revealed that the tumor capsule potentially affects intratumor spatial cluster continuity, transcriptome diversity and immune cell infiltration. Locally, we found that the bidirectional ligand-receptor interactions at the 100 μm wide cluster-cluster boundary contribute to maintaining intratumor architecture. Our study provides novel insights for diverse tumor ecosystem of PLCs and has potential benefits for cancer intervention.

## INTRODUCTION

Large-scale cancer genome projects have already revealed extensive intertumor and intratumor heterogeneities (*1, 2*). Recent single-cell omics studies, especially by single cell RNA-seq (scRNA-seq) technology, have greatly advanced our understandings of the tumor cell heterogeneities (*3*), tumor infiltrated immune cell sub-populations (*4*) and the features of tumor associated stromal cells (*5–8*) at single cell level. These studies provided many novel insights into tumor subtyping, tumor initiation and evolution, drug resistances and therapeutic targets. However, the scRNA-seq technology still has limitations. The most critical point is that the spatial and morphologic information is lost after the tissue dissociation into single cell suspension, making it hard to study the tumor spatial architecture. Although some *in situ* hybridization (ISH) based methods, such as MERFISH (*9*) and seqFISH (*10*) can obtain the spatial information, but they can only detect a few known target genes simultaneously.

The recently developed spatial transcriptomics (ST) technology (*11*) could overcome the above limitations. By positioning histological cryosections on arrayed reverse transcription primers with unique positional barcodes, ST provides high-quality genome-wide transcriptome data with intact two-dimensional positional information (*12*). It has been applied to analyzing the spatial heterogeneity of human primary breast cancer (*13*), melanoma (*14*), prostate cancer (*15*), pancreatic ductal adenocarcinomas (*16*) and human heart (*17*), etc. However, due to the relative lower resolution of former spatial transcriptomics method (maximal 1007 spots of 100 μm diameter and 200 μm interval) (*16*) and the lack of sequential comparison from adjacent normal to tumor inside region, the spatial architecture and heterogeneous tumor microenvironment (TME) have not been fully addressed.

Primary liver cancer (PLC) is the second most mortality tumors, of which hepatocellular carcinoma (HCC) and intrahepatic cholangiocarcinoma (ICC) are the two major histologic subtypes (*18*). Etiological and biological diversities, comprising chronic hepatitis virus infection, excess stress, drug-induced liver injury, aflatoxin B exposure, un-resolving inflammation and complicated TME, contribute to the high degree of intratumor heterogeneity of PLC (*19*). Till now, there are few effective non-surgical strategies for PLC, and lack of specific drug targets for effective therapeutic intervention (*20*). Only a limited proportion of HCC patients could benefit from existing tyrosine kinase inhibitor drugs (such as sorafenib and lenvatinib) and immune checkpoint inhibitors (*21–23*), largely resulting from both intertumor and intratumor heterogeneities (*24*). Till now, several efforts have been made to define the tumor heterogeneity and its clinical significance for liver cancer (*25–27*). For instance, scRNA-seq revealed that specific T cell subsets such as exhausted CD8^+^ T cells and regulatory T cells (Tregs) are preferentially enriched and clonally expanded in HCC (*4*); similarly, a VEGF/NOTCH-involved immunosuppressive onco-fetal TME was identified in HCC tumorigenesis (*28*). Moreover, our recent study also reported that the enrichment of CD4/CD8/PD1 triple-positive T cells in the tumor leading-edge region significantly indicates better prognosis (*29*), which reinforces that it is indispensable for comprehensive and accurate assessment of spatial heterogeneity for understanding the tumor cell community.

Here, we for the first time determined the spatial transcriptome architecture of 7 PLCs including total 84,823 tissue spots within 21 sections, and characterized the TME features including stromal and immune cell distribution, tumor cluster interaction. These findings provide novel insights for the complex ecosystem of liver cancer and have the potential to improve individualized cancer prevention and drug discovery.

## RESULTS AND DISCUSSION

### Exploration of Primary Liver Cancer Architecture with Spatial Transcriptomics

In order to comprehensively analyze the spatial heterogeneity of primary liver cancer (PLC), we collected 21 tissue specimens from 7 patients including five cases of hepatocellular carcinoma (HCC-1, 2, 3, 4, 5), one case of combined hepatocellular and cholangiocarcinoma (cHC-1), and one case of intrahepatic cholangiocarcinoma (ICC-1) and applied spatial transcriptomics (ST) sequencing via 10X Genomics Visium platform (Fig. 1A). For HCC-1/2/3/4 and cHC-1, we used three sequential sections (N: non-tumor section; L: leading-edge section; T: tumor section). For ICC-1, only L-section was collected due to massive necrosis inside the tumor. For HCC-2, an extra section from portal vein tumor thrombus (P-section) was collected. For HCC-5, the intact tumor nodule (diameter about 1 cm) was cut into four parts (designated as HCC-5A, B, C, D) to form a complete plane for ST analysis (Table S1). The bulk tissues for all the sections were also used for whole exome sequencing (WES) with peripheral blood mononuclear cell (PBMC) as control.

**Figure 1.**
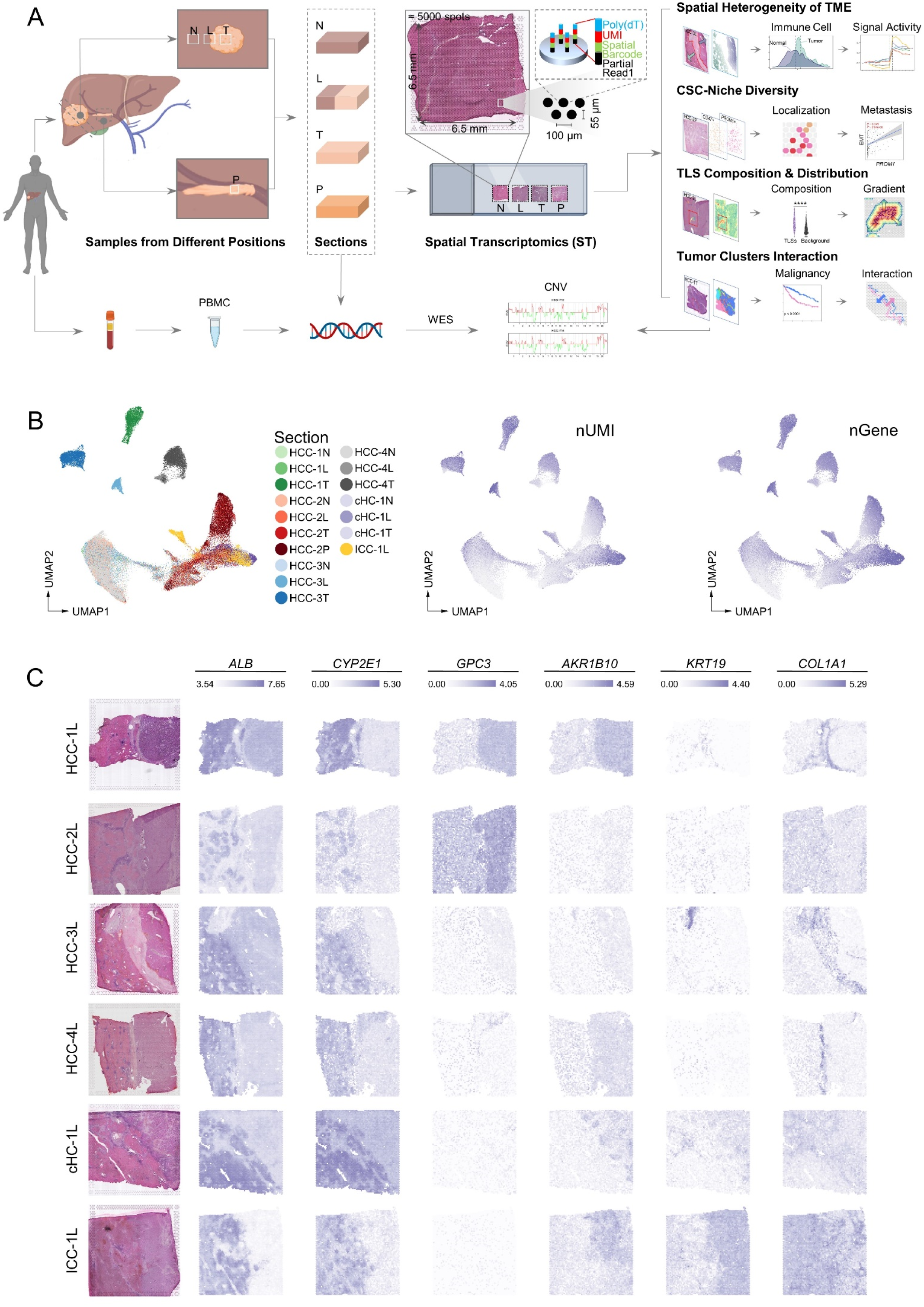
Exploration of primary liver cancer architecture with spatial transcriptomics. (**A**) Workflow of primary liver cancer samples collection, processing for ST and WES, and data analysis. N: Non-tumor area; L: Leading-edge area; T: Tumor area; P: Portal vein tumor thrombus area. (**B**) UMAP (Uniform manifold approximation and projection) plot of spots from all sections, colored by their sample source, the number of expressed UMIs (nUMI) and genes (nGene), respectively. HCC-1N represented the N-section of HCC-1. (**C**) H&E staining and the spatial feature plots of six marker genes of each L-section.

For the ST technology in this study, the diameter of spot reached 55 μm that captured approximately 8-20 cells according to H&E images (fig. S1A), and each section contained up to 5,000 spots in its capture area (6.5*6.5 mm^2^). Data showed the median sequencing depth of single spot at approximately 30,000 Unique Molecular Identifiers (UMIs) and 3,000 genes in this study (Table S2). Generally, the numbers of UMIs in tumor regions were larger than that in normal regions, consistent with previous studies (*30, 31*) (Fig. 1B and fig. S1B).

To verify whether the transcriptomic features are consistent with the histological information, we compared the H&E staining images with their counterpart ST data regarding the expression of several marker genes. Results confirmed that the regions defined by cell type marker genes’ expressions were highly consistent with the pathological images. Specifically, *ALB* and *CYP2E1* were highly expressed in normal regions; *GPC3* and *ARK1B10* in tumor regions; *KRT19* in cholangiocarcinoma regions; *COL1A1* in capsule and stromal regions (Fig. 1C).

### Different Patterns of PLC Spatial Heterogeneities

To characterize the spatial diversity of the PLCs, we combined the spots from different sections for each patient and performed clustering analyses (*30, 32, 33*). The distribution of the clusters was presented in both the UMAP projection space and tissue physical space. As shown in Figure 2A, we found that the clusters in HCC-1T, HCC-3T and HCC-4T had the characteristics of regional distribution whereas the clusters in HCC-2T and cHC-1T were intertwined. Cluster-5 in HCC-3L was a unique cluster that not appeared either in HCC-3N or HCC-3T.

**Figure 2.**
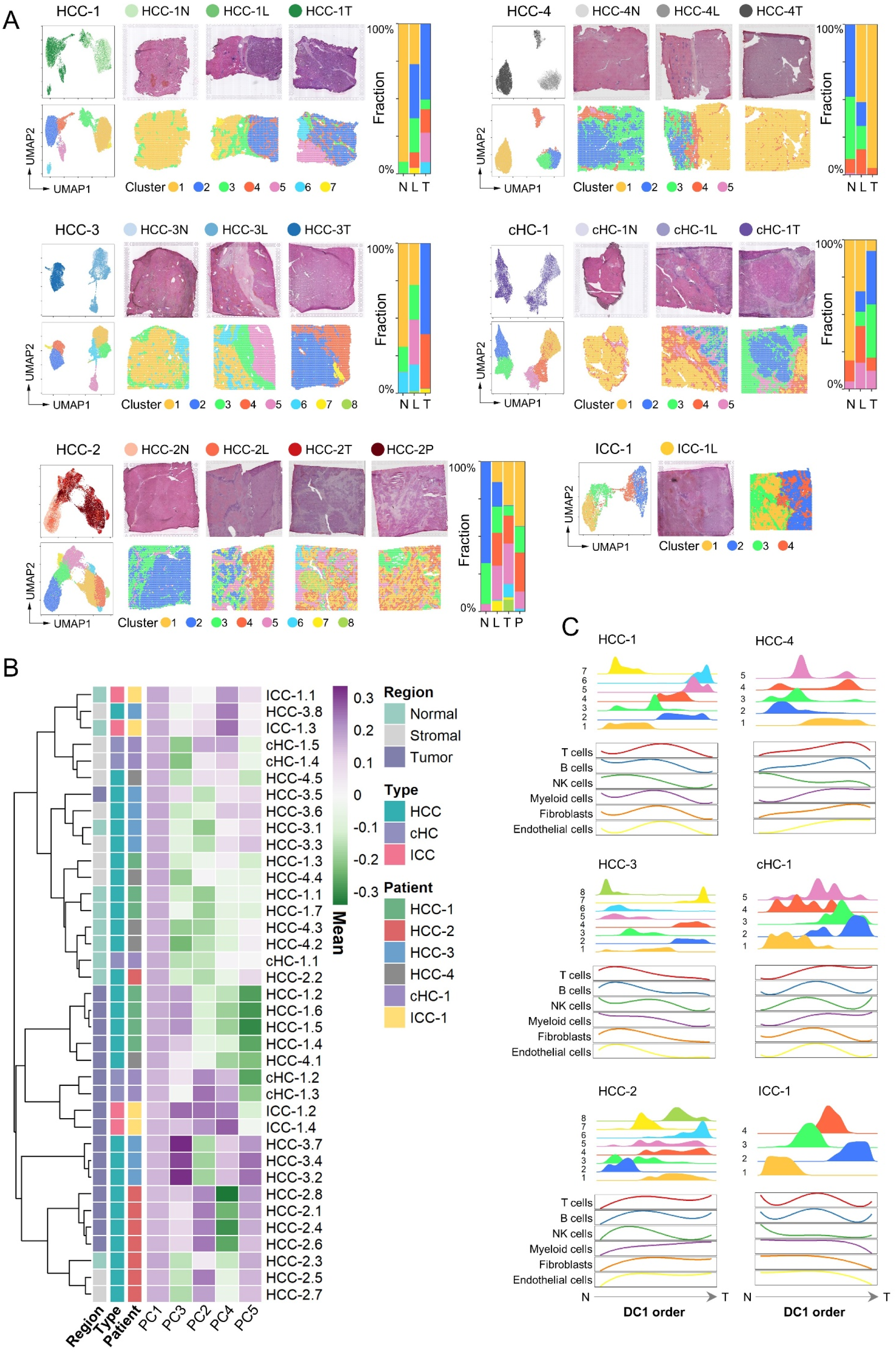
Different patterns of PLC spatial heterogeneities. (**A**) For each patient: left, UMAP of the spots colored by their section sources and cluster identities, respectively; middle, H&E staining and the spatial cluster distribution of each section; right, the fraction of clusters in each section. (**B**) Similarity comparison of the clusters across different patients. The clusters’ tissue regions, histopathological types, and patient information were annotated on the left. HCC-1.1 represented the cluster-1 of HCC-1. (**C**) Distribution of the clusters and the main stromal and immune cell scores along the direction from normal (N) to tumor (T). This direction was estimated by the first components of the spots’ diffusion map (DC1), which were generally ordered from normal to tumor.

To examine the subtypes at cluster level across different patients, we performed hierarchical clustering and diffusion map analysis (*34*), and found that the clusters from the same type of regions were more similar in general (Fig. 2B and fig. S2A, left panel). In the diffusion map, a three-branch structure was formed: the normal and tumor region clusters were projected into the branches and the stromal region clusters were on the junction (fig. S2A, right panel).

Next, we tried to explore the spatial distribution of different cell types. Considering that each spot may contain more than one cell, we proposed a signature-based strategy to score the enrichment of different cell types in each spot (*35*) (Table S3). Notably, the fibroblast and endothelial cell scores were significantly higher in stromal regions, and the immune cell scores in the different clusters of tumor regions exhibited high degree of diversity (fig. S2B). As a validation, we performed the multimodal intersection analysis (MIA) (*16*) by integrating our ST data with a liver cancer single cell data set (*19*), and got the similar results (fig. S2C and D). Then, to analyze the composition changes of cell types from the outside to the inside of tumor, we used diffusion map to project the spots into a one-dimensional pseudo order (the first diffusion component, DC1) (*34*), which can be seen as generally from normal to tumor regions by comparing the distribution of clusters in this order. By fitting the variation curves of cell type scores (including T, B, NK, myeloid, endothelial cells and fibroblasts), we found that the variation patterns of T, B, endothelial cells and fibroblasts were similar with each other in five cases except cHC-1, whereas that of NK and myeloid cells was highly variable (Fig. 2C).

### Microenvironment Characteristics in Leading-edge Area

As seen in Figure 2A, the tumor clusters’ spatial distribution presented two distinct patterns. One was block-like with clear boundary between clusters (e.g., HCC-1T, HCC-3T), while the other was discontinuous and mixed (e.g., HCC-2T). To measure this characteristic quantitatively, we introduced a metric named “spatial continuity degree”, which was calculated by comparing the consistency of cluster identity between each spot and its neighbors. Together with another metric, “transcriptome diversity degree”, which measured the global transcriptomic heterogeneity of tumor regions in each section, we quantitatively found that the tumor regions of L&T-sections in HCC-1, HCC-3 and HCC-4 patients had higher spatial continuity and lower transcriptome diversity (Fig. 3A).

**Figure 3.**
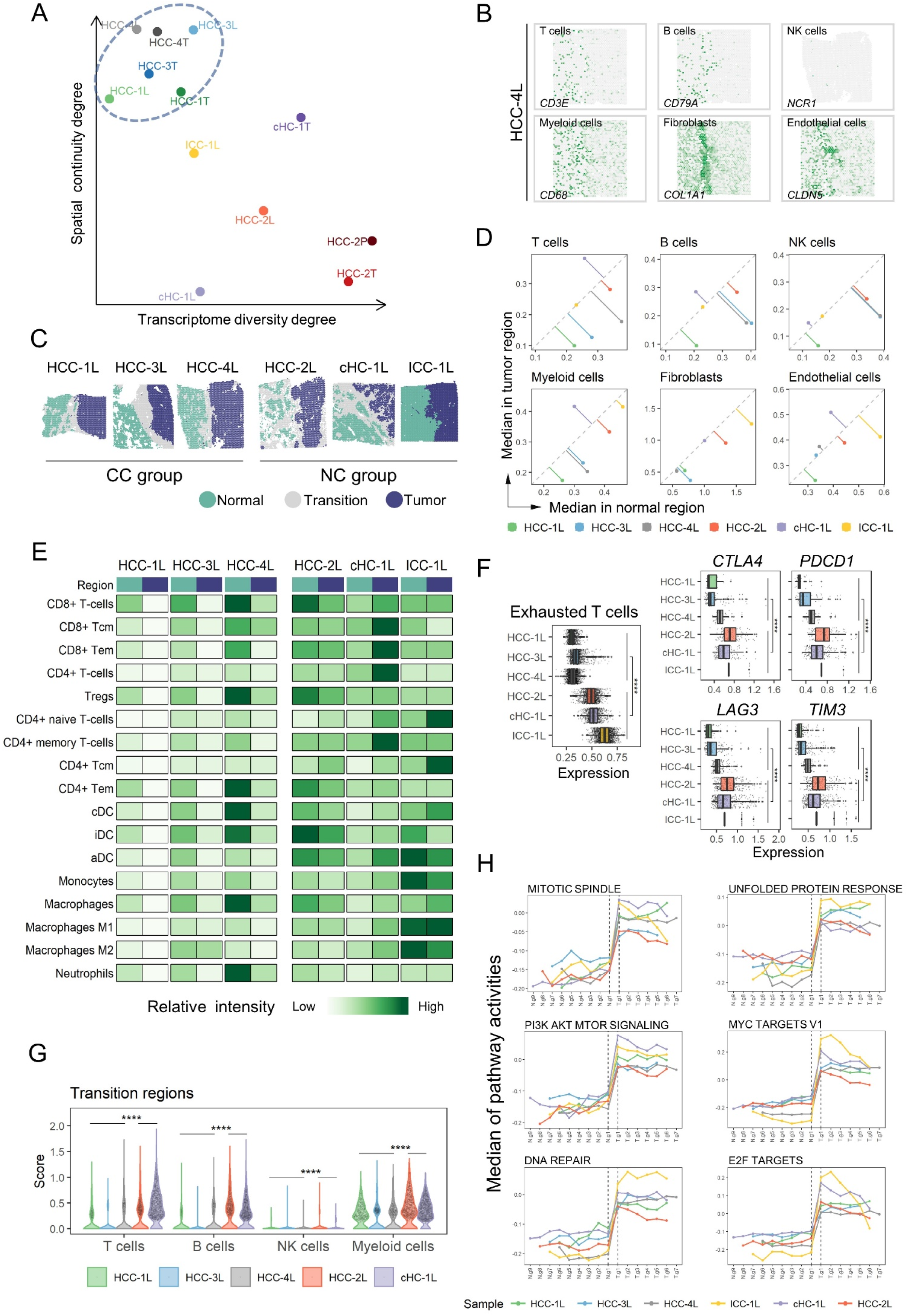
Microenvironment characteristics in leading-edge area. (**A**) The transcriptome diversity degree and spatial continuity degree of tumor regions in L/T/P-sections. (**B**) The spatial feature plots of six marker genes of stromal and immune cell types in HCC-4L. (**C**) The distribution of normal, tumor and transition regions in the L-sections, and the grouping results of the CC (complete capsules) group and NC (non or discontinued capsules) group. (**D**) Comparison of the median of stromal and immune cell type scores between the normal (x-axis) and the tumor (y-axis) regions in each L-section. (**E**) Comparison of the relative intensity (each row shared a color scale, while different rows didn’t) of stromal and immune cell subtype scores between the normal and tumor regions in each L-section. (**F**) Comparison of the expression levels of exhausted T cell signature, *CTLA4*, *PDCD1*, *LAG3* and *TIM3* between the CC and NC groups. Two-sides Wilcoxon rank-sum tests on the CC and NC groups were used to analyzed the significance of their differences. **** p<0.0001. (**G**) Comparison of the immune cell scores in the “transition regions” between the CC and NC groups. One-sided Wilcoxon rank-sum tests (the CC group was less than NC group) were used to calculated the statistical significance. ****, p<0.0001. (**H**) The changes of hallmark pathways’ activities along the gradient divisions on the both sides of transition region. Each dot indicated the median of the pathway activity in the corresponding area.

To further explore the common features of those three samples with higher spatial continuity and lower transcriptome diversity (HCC-1L, HCC-3L, and HCC-4L), we investigated their clinicopathological features (Table S1) and found that all of them had complete capsules. Whereas the capsules of cHC-1L and HCC-2L were pathologically incomplete, and ICC-1L had no capsule at the border of the tumor nodule. It was observed that the expression of stromal and immune cell markers displayed sharply reduction across the capsule from the normal side to tumor side in HCC-4L, indicating the capsule may affect the stromal and immune cell distribution (Fig. 3B). We thus defined the capsules of HCC-1L, HCC-3L, and HCC-4L and stromal cell clusters of cHC-1L and HCC-2L as the “transition area” in L-sections, and investigated the spatial characteristics of the “normal region” and “tumor region” on the two sides of the transition area. Meanwhile, HCC-1L, HCC-3L and HCC-4L were named as “complete capsules (CC)” group, while HCC-2L, cHC-1L and ICC-1L as “non or discontinued capsules (NC)” group (Fig. 3C). As shown in Figure 3D, the scores of T, B and myeloid cells were much higher in normal area than in tumor area in CC group instead of NC group. The fibroblast and endothelial cell subtype scores were lower in both tumor and normal regions of CC group compared with NC group. CD8^+^ Tem (effective memory T cells), Tregs (regulatory T cells), CD4^+^ memory T cells, CD4^+^ Tem, cDC (conventional dendritic cell), monocytes, macrophages M1 and neutrophils were found significantly enriched in normal regions in CC group (Fig. 3E). By comparing the function of T cells in tumor region between two groups, we found that exhausted T cells were dramatically increased in NC group in coupled with the increased expression of *PDCD1*, *CTLA4*, *LAG3* and *TIM3* (Fig. 3F). In addition, the scores of T, B, NK and myeloid cells in the transition regions were significantly lower in the CC group (Fig. 3G).

Since swelling and invasive growth is the biological characteristic of tumor cells, we wondered whether the signaling pathway activities had gradient-like changes in the direction perpendicular to the boundary on both sides of the capsule. To address this question, we divided the normal and tumor regions of L-sections into continuous zones parallel to the shape of the dividing line at intervals of 5 spots (fig. S3A) and analyzed the activities of hallmark pathways by gene set variation analysis (GSVA) (*36*). Generally, the majority of hallmark pathways did not show gradient in either normal or tumor region and few consistent patterns were observed in different patients (fig. S3B). Individually, the ICC section ICC-1L showed several distinct changes: hypoxia-associated signals showed sudden decrease from N.g1 to T.g1 zone, whereas increased dramatically from T.g1 to T.g6 zone, and inflammatory response and interferon response (alpha and gamma) pathways were observed gradually reduced from T.g1 to T.g6 zone (fig. S3B). It should be noted that six classical tumor-associated pathways (including Phosphoinositide 3-kinase, MYC, mitotic spindle, unfolded protein response, E2F and DNA repair) exhibited sudden increase from N.g1 to T.g1 zone regardless of the width of the transition area or the presence of capsule (Fig. 3H).

Taken together, our study here indicates that the integrity of the capsule is closely associated with the spatial heterogeneity of tumor cells and the distribution of their surrounding stromal and immune cells, but has few effects on the activities of hallmark pathways in either normal or tumor regions.

### Intratumor Heterogeneity in PLCs

To investigate the interior heterogeneities of tumor regions in both L&T-sections, we calculated hallmark pathways’ activities for the spots in tumor regions. By performing hierarchical clustering on the averaged pathway activities in each tumor cluster, two modules were identified (*37*)(Fig. 4A). Module-1 showed high activities of cell cycle and metabolism-related pathways (e.g., MYC targets v1, G2M checkpoint, E2F targets, cholesterol homeostasis, bile acid and fatty acid metabolism, etc.), while Module-2 had much higher activities in the inflammation, angiogenesis and epithelial-mesenchymal transition (EMT) pathways.

**Figure 4.**
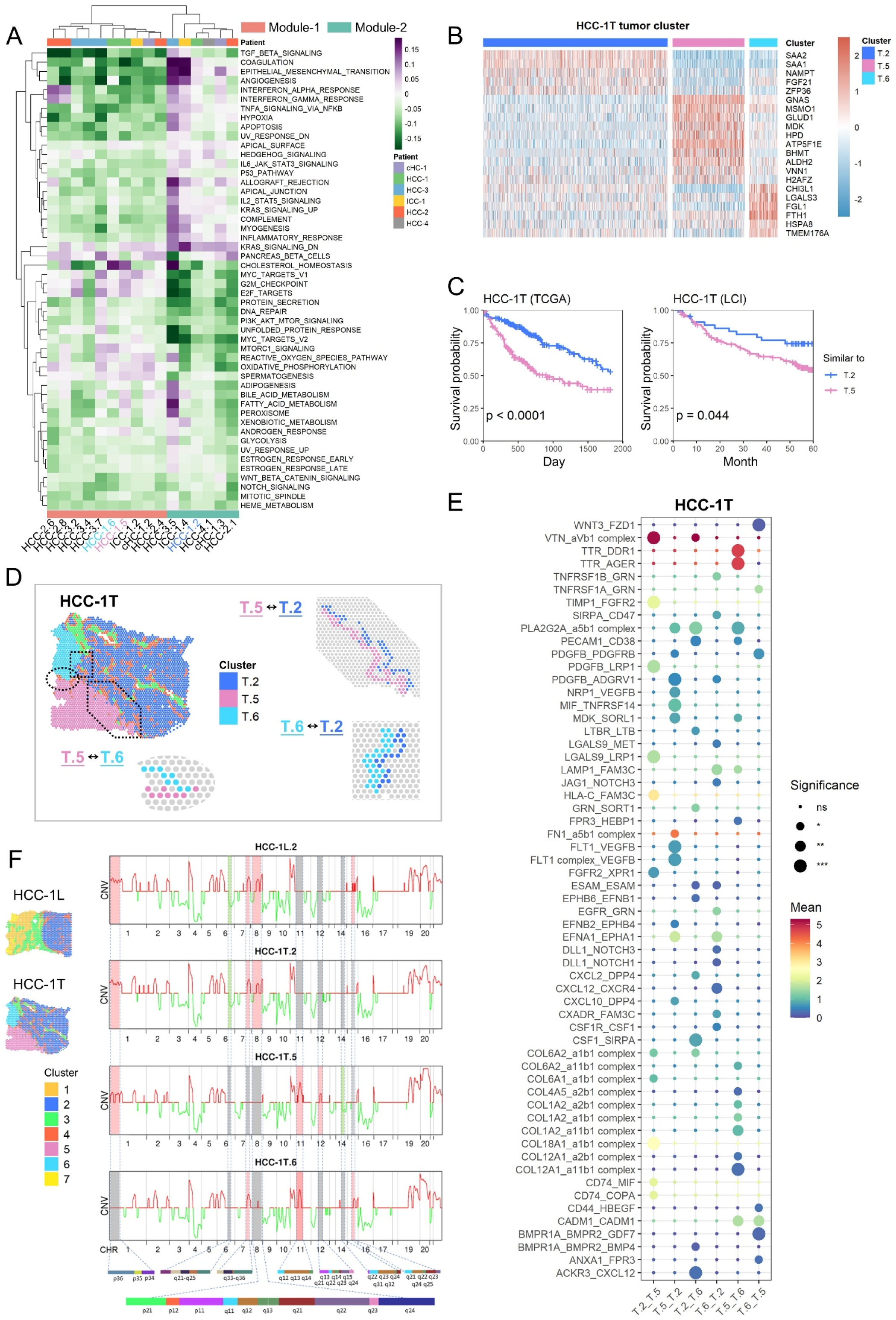
Intratumor heterogeneity in PLCs. (**A**) Clustered heatmap of tumor clusters’ hallmark pathway average activities. The tumor clusters were grouped into two functional modules. HCC-1.2 represented the cluster-2 of HCC-1. (**B**) Expression profiles of some differential expression genes of the clusters-2/5/6 in HCC-1T. T.2 represented the cluster-2 in HCC-1T. (**C**) The survival curves of two groups of patients in TCGA and LCI cohorts to compare the relative malignancy of ST tumor cluster pairs (cluster-2 vs 5 in HCC-1T). These two groups were divided according to which ST tumor cluster the bulk samples were more similar to at expression level. Log rank test was used to measure the statistical significance of their relative malignancy degrees. (**D**) The definition of the boundary areas to study the interaction between two neighbor tumor clusters in HCC-1T. The regions with 4 spots wide along the boundary lines in each cluster were selected and the spots of stromal clusters were excluded. (**E**) Bubble heatmap showing the mean interaction strength between the neighbor clusters at the boundaries for ligand-receptor pairs. Dot size indicated the statistical significances by permutation test. Dot color indicated the mean interaction strength levels. HCC-1T.2 represented the cluster-2 in HCC-1T. (**F**) The averaged CNV profiles for each tumor cluster in HCC-1, inferred from spatial transcriptomes. The color of the lines indicated the amplifications (red) and deletions (green). The differences between clusters were highlighted by background colors (red, green, and grey) and their detailed chromosome band labels were also presented. HCC-1L.2 represented the cluster-2 in HCC-1L.

The tumor clusters of HCC-1 belonged to two different modules, T.2 (represented cluster-2 in HCC-1T) to Module-2 and T.5&6 to Module-1, showing distinct gene expression patterns (Fig. 4B and Table S4). By comparing these three tumor clusters with the samples of two HCC bulk transcriptome datasets (The Cancer Genome Atlas [TCGA] cohort and the Liver Cancer Institute [LCI] cohort) (*38, 39*), we found that bulk samples more similar to T.2 had better prognosis than those similar to T.5&6 (Fig. 4C and fig. S4A). Moreover, bulk samples more similar to T.5 showed even worse outcome than that similar to T.6 in TCGA cohort (fig. S4B), implying the up-regulated creatine, tyrosine, ethanol and retinol metabolism pathways in T.5 may enhance cell malignant behaviors, and new strategy targeting those metabolism pathways could be tested for HCC intervention.

To investigate the communication and interaction between tumor clusters in HCC-1, the interface regions of clusters were selected with the range of 4 spots wide (2 spots wide for each cluster; the spots in stromal regions were excluded) (*40, 41*) (Fig. 4D). It was found that the enriched gene-pairs including NRP1-VEGFB (*42, 43*), FLT1-VEGFB (*44*), EFNB2-EPHB4 (*45*), MDK-SORL1 (*46*) and EFNA1-EPHA1 (*47*) may contribute to T.5 induced cell angiogenesis/proliferation/migration in T.2. In turn, T.2 may help to maintain the metabolic activity in T.5 through LGALS9-LRP1, PDGFB-LRP1 (*48*), et al. SORL1, sortilin-related receptor 1, has been reported involving endosomal trafficking and oncogenic fitness, which might help to induce drug resistance and tumor cell growth (*46*) in T.2. Meanwhile, LRP1, low-density lipoprotein receptor-related protein 1, is a multifunctional receptor involved in endocytosis and metabolism homeostasis (*49*), the potential role of which should be explored in future. Similar patterns also existed in HCC-1T.2 & HCC-1T.6 (HCC-1T.2 represented cluster-2 in HCC-1T) and HCC-1T.5 & HCC-1T.6 interactions (Fig. 4E). Together, our data here suggest that those molecules high-expressed in each cluster could mediate reciprocal communications and might be used as potential targets to disrupt tumor cell communities for clinical treatment.

To study possible genomic drivers of different tumor clusters spatially, we inferred the copy number variations (CNVs) from ST data and WES data of matched bulk tissues (*33, 50, 51*). It can be seen that most CNVs inferred from ST data were consistent with WES bulk data, which suggests that the inferred CNVs from ST data were reliable. Further, ST data generated more subtle CNV heterogeneities across different tumor clusters. For example, in HCC-1: 1) HCC-1L.2 had exactly the same CNV pattern as in HCC-1T.2, verifying that the same clusters across different sections had identical CNV characteristics; 2) the observation that most CNV regions were shared across T.2/5/6 in HCC-1 suggested that those three clusters might be derived from the same clone; and 3) the gain of chromosome 11q13 was found in HCC-1T.5&6, but not in HCC-1L.2/HCC-1T.2, whereas chromosome 8q13 amplifications were in HCC-1L.2/HCC-1T.2, but not in HCC-1T.5&6 (*52, 53*) (Fig. 4F and figs. S5A and S5C). In contrast, for HCC-3, CNV regions in L.5 were obviously different from T.2/4/7, indicating the distinct origin of T.5 (figs. S5B and S5D).

Taken together, these results suggest that the spatial intratumor heterogeneities exist widely. The different clusters within certain tumor nodule have diversified pathway activities and distinct origins. The reciprocal communications across different clusters might be essential for tumor ecosystem and evolution.

## DISCUSSION

In this study, genome-wide heterogeneity transcriptomes of seven primary liver cancers were measured with a 55 μm spatial resolution for the first time. By conducting sequential comparison from normal to leading-edge to tumor regions, we found that the tumor capsule potentially affects intratumor spatial cluster continuity, transcriptome diversity, immune cell infiltration and cancer hallmark pathway activities. Meanwhile, cell-cell interactions in the 100 μm wide tumor cluster-cluster boundary were comprehensively analyzed.

The tumor microenvironment (TME) comprises tumor, stromal and immune cells, extracellular matrix, and signaling molecules (*54, 55*). The spatially and temporally dynamic variations in TME are considered as the key factors for tumor heterogeneity (*56*). To our knowledge, this is the first study to sequentially analyze the genome-wide TME characteristics from normal to leading-edge to tumor regions, which provide us the opportunity to investigate both the global and local variation tendency of difference cell populations. These results show that the capsules, mainly consisting of fibroblasts and endothelial cells, can act as a barrier preventing the infiltration immune cells, which is supported by a few previous studies (*57*). More importantly, we found that the absence of capsules in L-sections could lead to both lower spatial continuity and higher transcriptome diversity in tumor. Together with the observation that the relatively small TLS spots in L&T-sections in HCC-1/3/4, our data here reinforce the key role of capsule for both TME architecture and intratumor heterogeneity.

The tumor heterogeneity includes interpatient heterogeneity, intertumor heterogeneity (different tumor nodules within the same patient) and intratumor heterogeneity (different regions in the identical tumor nodule) (*58*). Globally, the spatial distribution of clusters within tumors has two distinct patterns: regional distribution and intertwined distribution. The regionally distributed clusters tend to have higher spatial continuity and lower transcriptome diversity. By comparing the inferred CNVs from ST data with the CNVs from matched bulk WES data, most CNVs were consistent, reinforcing that it is practical to infer CNV by ST data. It should be noted that in comparison with cluster-2/4/7 in HCC-3T, cluster-5 displayed distinct CNV pattern and significant lower level of genomic disorder. Together with the pathological stage that cluster-5 was focal nodular hyperplasia (FNH, the pre-malignant nodule) instead of neoplasia, these results suggest the distinct and initial evolution trajectory of cluster-5 instead of cluster-2/4/7 in HCC-3 (Fig. 4). By establishing a complete spatial transcriptome of an intact tumor nodule (diameter around 1 cm) collected from an early stage HCC patient, we found that the heterogeneous tumor clusters have already existed in such small tumor nodule, implying the formation of diverse populations during tumorigenesis. Regarding the changes of signal activities, we uncovered widely different trends of hallmark pathway activities in all directions from inside to outside of tumor, which might result from the interplay with other cluster cells and multiple stresses locally. This complex heterogeneity of HCC explains the reason why the current drug treatment for HCC is usually ineffective, and also suggests that a comprehensive detection of the genetic characteristics of different parts of the tumor may make a breakthrough in immunotherapy.

The emerging spatial transcriptomics technology could largely solve the shortcomings of missing cell spatial location information by single-cell sequencing, and play an important role in many research fields. However, the major limitations of ST are still its resolution: 1) although the diameter of each spot in ST section in our present study has reached 55 μm instead of 100 μm in former version (8-20 cells vs 100-200 cells each spot) (*16*), it is still unable to provide the comparable accuracy at single-cell scale; 2) ST method can only provide the transcript information of cells within spots, whereas the information of interval spaces between two spots is missed. To deal with the first issue, we applied the cell type signature-based strategy to calculate the enrichment of TME cell types in each spot, which achieved similar result as previous developed multimodal intersection analysis (MIA) method did (figs. S2B-D). For the second issue, the improved technology with shorter interval distance and advanced analytical method need to be developed in future.

Tumor heterogeneity is the major obstacle for liver cancer diagnosis and therapy. Our study presents the first genome-wide spatial transcriptome map of three major liver cancer subtypes. Extensive global and local intratumor heterogeneities of tumors and TMEs have been found. Also, the tumor clusters from different patients show distinct spatial patterns and transcriptomic diversities. These findings provide meaningful insights to find new drug targets and develop novel therapeutic strategies.

## MATERIALS AND METHODS

### Human primary liver cancer (PLC) samples and blood

The adjacent normal and tumor (PLC) tissues were collected under a protocol approved by the Ethics Committee of Eastern Hepatobiliary Surgery Hospital (EHBH) (EHBHKY2018-1-001). Individuals donating fresh surgical tissue provided informed consent. All diagnoses were verified by histological review by a board-certified pathologist. The samples were delivered within MACS Tissue Storage Solution (Miltenyi, Cat#: 130-100-008). For tumors of adequate size, small fragments of each tumor were snap frozen in optimum cutting temperature (OCT) compound (SAKURA, Cat#: 4583) and stored at −80 °C until use. Peripheral venous blood (5 ml) was collected in an anticoagulation tube (BD Vacutainer, Cat#: 367525).

### Collection and preparation of primary liver cancer (PLC) tissue

The surgically resected PLC tissue was immediately submerged in MACS Tissue Storage Solution and sent to the laboratory for processing as soon as possible. Then, PLC tissue was gently washed with cold PBS (GIBCO, Cat#: 20012-043) and cut into about 6.5 mm^3^ pieces (bulks) according to the experimental design. Also, the same pieces immediately adjacent to them were directly frozen at -80°C forwhole exome sequencing (WES). The tissue bulks were placed in OCT-filled mold and snap frozen in isopentane and liquid nitrogen. Cryosections were stored at −80 °C until use.

### Peripheral blood mononuclear cell (PBMC) preparation

Fresh anticoagulated blood was centrifuged (500 g, 5 min), and then the upper layer of plasma was removed. After adding 10 ml PBS and mixing thoroughly, the mixed solution was slowly added to the surface of Ficoll solution (GE Healthcare, Cat#: 17544203). After centrifugation (800 g, 20 min, room temperature, brake off), the white mononuclear cell layer in the middle of the solution was taken out and mixed with 10 ml PBS. After centrifugation again (500 g, 5 min), the solution was discarded. Finally, the sediment at the bottom was resuspended in 200 μl cell freezing medium (10% dimethyl sulphoxide + fetal bovine serum) and stored at -80°C.

### Spatial transcriptome sequencing

This experiment is based on the Visium Technology Platform of 10X Genomics company. The reagents and consumables in the experiment are provided by this platform, and the specific product numbers can be found at https://www.10xgenomics.com/products/spatial-gene-expression.

### Slide preparation

The Visium Spatial Gene Expression Slide (from Visium Spatial Gene Expression Slide Kit, 10X Genomics, PN-1000185) includes 4 capture areas (6.5*6.5 mm^2^), each defined by a fiducial frame (fiducial frame + capture area is 8*8 mm^2^). The capture area has ∼5,000 gene expression spots, each spot with primers that include: Illumina TruSeq Read 1 (partial read 1 sequencing primer); 16 nt Spatial Barcode (all primers in a specific spot share the same Spatial Barcode); 12 nt unique molecular identifier (UMI); 30 nt poly (dT) sequence (captures poly-adenylated mRNA for cDNA synthesis).

### RNA integrity number (RIN)

We use RNeasy Mini Kit (Qiagen, Cat#: 74104) to test the integrity of RNA. After taking 10 slices of 10 mm thickness cryosections, RNA was extracted and analyzed by RNeasy Mini Kit immediately. RIN≥7 is qualified.

### Optimization of the permeabilization time

Prior to using a new tissue for generating Visium Spatial Gene Expression libraries, the permeabilization time was optimized. Briefly, the Visium Spatial Tissue Optimization workflow included placing tissue sections on 7-capture areas on a Visium Tissue Optimization slide (from Visium Spatial Gene Expression Reagent Kit, 10X Genomics, PN-1000186). The sections were fixed, stained, and then permeabilized for different times. The mRNA released during permeabilization binds to oligonucleotides on the capture areas. Fluorescent cDNA was synthesized on the slide and imaged. The permeabilization time that results in maximum fluorescence signal with the lowest signal diffusion was optimal. If the signal was the same at two time points, the longer permeabilization time was considered optimal. Once optimal conditions had been established, each cryosection was cut at 10 mm thickness onto Visium Slide (from Visium Slide Kit), and processed immediately. In this study, the permeabilization time ranges from 6 to 24 min depending on the samples.

### Tissue fixation, staining and imaging

Tissue sections on the Visium Slide (from Visium Slide Kit) were fixed using methanol (Millipore Sigma) by incubating 30 min at -20°C. For tissue staining, sections were incubated in isopropanol (Millipore Sigma) for 1 min, in Hematoxylin (Agilent) for 7 min, in Bluing Buffer (Agilent) for 2 min and in Eosin Mix (Millipore Sigma) for 1 min at room temperature. Lastly, slides were incubated for 5 min at 37°C in the Thermocycler Adaptor (10X Genomics, PN-3000380). The slides were washed in ultrapure water after each staining steps. Then, the stained tissue sections are imaged.

### Tissue permeabilization and reverse transcription

For tissue permeabilization, the slides were first placed in the Slide Cassette (from the Visium Slide kit) for the optimal permeabilization time. A Permeabilization Enzyme (from the Visium Reagent kit) was used for permeabilizing the tissue sections on the slide for incubating for the pre-determined permeabilization time. The poly-adenylated mRNA released from the overlying cells was captured by the primers on the spots. After washing by 0.1*SSC (Millipore Sigma), RT Master Mix (provided in Visium Reagent kit) containing reverse transcription reagents was added to the permeabilized tissue sections in the Thermocycler Adaptor. Incubation with the reagents produces spatially barcoded full-length cDNA from polyadenylated mRNA on the slide.

### Second strand synthesis and denaturation

After removing RT Master Mix (provided in Visium Reagent kit) from the wells, sections were incubated in 0.08 M KOH for 5 min and washed by Buffer EB (Qiagen). Then, Second Strand Mix (provided in Visium Reagent kit) was added to the tissue sections on the slide to initiate second strand synthesis on the Thermocycler Adaptor. This is followed by denaturation and transfer of the cDNA from each Capture Area to a corresponding tube for amplification and library construction. The slides were washed by Buffer EB and incubated in 0.08M KOH for 5 min. Then, samples from each well were transferred to a corresponding tube containing Tris-HCl (1 M, pH 7.0) in 8-tube strip for amplification and library construction.

### cDNA amplification and QC

1 μl sample from Denaturation was transferred to the qPCR plate well containing the qPCR Mix (Nuclease-free water + KAPA SYBR FAST qPCR Master Mix (KAPA Biosystems) + cDNA Primers (from Visium Reagent kit)). The Cq Value for each sample was recorded after qPCR implemented. For cDNA amplification, cDNA Amplification Mix (from Visium Reagent kit) was added to the remaining sample from Denaturation. Then, the product was incubated in Thermocycler Adaptor for a cycle. For cDNA Cleanup–SPRIselect, 60 μl SPRIselect reagent (Beckman Coulter) was added to each sample and incubated for 5 min at room temperature. The sample was repeatedly adsorbed by the magnet•High, washed with ethanol (Millipore Sigma) and Buffer EB, and transferred to a new tube strip. Then, run 1 μl of sample on an Agilent Bioanalyzer High Sensitivity chip (Agilent, Cat#: 50674626) for cDNA QC & Quantification.

### Visium spatial gene expression library construction

Enzymatic fragmentation and size selection were used to optimize the cDNA amplicon size. P5, P7, i7 and i5 sample indexes, and TruSeq Read 2 (read 2 primer sequence) were added via End Repair, A-tailing, Adaptor Ligation, and PCR. The final libraries contain the P5 and P7 primers used in Illumina amplification. Library construction was performed with Library Construction Kit (10X Genomics, Cat#: PN-1000190).

### Fragmentation, end repair & A-tailing

Only 10 µl purified cDNA sample from cDNA Cleanup was transferred to a tube strip. Buffer BE and Fragmentation Mix (from Library Construction kit) were added to each sample, and Fragmentation was performed in thermal cycler. Post Fragmentation, 30 µl SPRIselect reagent (0.6X) was added to each sample and incubated for 5 min at room temperature. The sample was repeatedly adsorbed by the magnet•High, washed with ethanol and Buffer EB, and transferred to a new tube strip.

### Adaptor ligation

50 μl Adaptor Ligation Mix (from Library Construction kit) was added to each 50 μl sample and incubated in a thermal cycler.

### Post ligation cleanup–SPRIselect

80 µl SPRIselect reagent (0.8X) was added to each sample and incubated for 5 min at room temperature. The sample was repeatedly adsorbed by the magnet•High, washed with ethanol and Buffer EB, and transferred to a new tube strip.

### Sample index PCR

50 µl Amp Mix (from Library Construction kit) and 20 µl of an individual Dual Index TT Set A (10X Genomics, Cat#: PN-1000215) was added to each 30 μl sample and incubated in a thermal cycler.

### Post sample index PCR double-sided size selection – SPRIselect

60 µl SPRIselect reagent (0.6X) was added to each sample and incubated for 5 min at room temperature. After adsorbing by the magnet•High, 150 µl supernatant was transferred to a new tube strip. Then, 20 µl SPRIselect reagent (0.8X) was added to each sample and incubated for 5 min at room temperature. After adsorbing by the magnet•High and supernatant removed, samples were washed with ethanol and Buffer EB, and transferred to a new tube strip.

### Post library construction QC

Run 1 µl of sample (1:10 dilution) on an Agilent Bioanalyzer High Sensitivity chip.

### Sequencing

A Visium Spatial Gene Expression library comprises standard Illumina paired-end constructs which begin and end with P5 and P7. The 16 bp Spatial Barcode and 12 bp UMI are encoded in Read 1, while Read 2 is used to sequence the cDNA fragment. i7 and i5 sample index sequences are incorporated. TruSeq Read 1 and TruSeq Read 2 are standard Illumina sequencing primer sites used in paired-end sequencing.

### Space ranger

The Visium spatial RNA-seq output and brightfield and fluorescence microscope images were analyzed by Space Ranger (version 1.1.0) in order to detect tissue, align reads, generate feature-spot matrices, perform clustering and gene expression analysis, and place spots in spatial context on the slide image. These pipelines combined Visium-specific algorithms with the widely used RNA-seq aligner STAR.

### Whole exon sequencing (WES)

Whole exome sequencing was performed per standard protocols using the Next-Generation Sequencing platform of TWIST bioscience company whose details can be found at https://www.twistbioscience.com/products/ngs. Briefly, for DNA extraction, snap frozen fresh biopsy and matched whole blood samples were processed using QIAamp DNA Mini Kit (Qiagen, Cat#: 51304), QIAamp DNA Blood Maxi Kit (Qiagen, Cat#: 51194) according to the manufacturer’s instructions, and quantified using the Qubit dsDNA BR Assay Kit (Thermo Fisher, Cat#: Q32853). Libraries were generated with Twist Library Preparation EF Kit 1 (TWIST, Cat#: 100572), Twist CD Index Adapter Set (TWIST, Cat#: 100577) and Twist Library Preparation Kit 2 (TWIST, Cat#: 100573). Subsequently, hybridization and capture were performed using the Twist Fast Hybridization and Wash Kit (TWIST, Cat#: 101175) and Twist Binding and Purification Bead (TWIST, Cat#: 100983), respectively. After capture, Libraries were amplified by PCR. Then, purified libraries were validated and quantified using an Agilent Bioanalyzer High Sensitivity DNA Kit (Agilent, Cat#: 50674626) and a Qubit dsDNA High Sensitivity Quantitation Assay (Thermo Fisher Scientific, Cat#: Q32854). Finally, the enriched libraries were sequenced on the Nova6000 instrument of Illumina platform (Illumina).

## Statistical analysis

### Spatial transcriptomics data processing

For the gene-spot matrixes generated by Space Ranger, some routine statistical analyses were performed firstly, including calculating the number of the detected UMIs (nUMI), and genes (nGene) in each spot. Based on them, the basic quality controls (QC) were applied on the data. In detail, the spots with extremely low nUMI or nGene (outliers), and the spots isolated from the main tissue sections were removed. The genes expressed in less than 3 spots, and mitochondrial, ribosomal genes were filtered.

After QC, we used the R package harmony (v1.0) (*30*) to integrate the expression data from different sections of each patient, and used the Seurat package (v3.1.5) (*32*) to perform the basic downstream analysis and visualization (*33*). In detail, we firstly combined the expression matrixes of each patient’s all sections, and performed normalization, log-transformation, centering and scaling on them. Next, we identified 2,000 highly variable genes according to their expression means and variances. Based on them, principal components analysis (PCA) was performed to project the spots into a low-dimensional space, which was defined by the first 20 principal components (PCs). Then, by setting the section source as the batch factor and using the “RunHarmony” function, we iteratively corrected the spots’ low-dimensional PC representation to reduce the of impact of batch effect. After this step, the corrected PC matrixes were used to perform unsupervised shared-nearest-neighbor-based clustering and UMAP (uniform manifold approximation and projection) visualization analysis further. And to compare the clusters at gene level, we identified differentially expressed genes of the all or selected clusters by using fold-change analysis and Wilcoxon Rank Sum test with Bonferroni correction.

### Cluster similarity analysis

For the clusters from different patients, we represented them by their spots’ average expression profiles (the log-transformed normalization values). To reduce the impact of extreme values, we excluded some outlier spots in advance, whose first three PC values beyond the range of the mean*±*3*standard deviation of the cluster they belonged to. Moreover, only the genes with the mean above 0.1 and the variance above 0.05 across all the cluster expression vectors were retained for the downstream comparison analyses.

To measure the clusters’ similarities across patients, we preformed two types of analyses, hierarchical clustering and low-dimensional projection. In detail, we firstly applied PCA on the centered and scaled clusters’ average expression profiles, and used the first five PCs to perform hierarchical clustering (Fig. 2B). Besides, the diffusion map was used to project clusters of different patients into a two-dimension space (the first two diffusion components) based on the package destiny (*34*) with default parameter setting (fig. S2A). For convenience of comparison, we annotated each cluster with a region label (normal, stromal, or tumor), which was decided by integrating the information of the cluster’s marker genes and H&E staining images.

### Cell type scoring by a signature-based strategy

At the current Visium ST resolution, each spot may contain approximately 8-20 cells, so that we couldn’t assign a certain cell type for each spot. Considering this, to compare the distribution of cell types across the tissue sections, we proposed a signature-based strategy to score the cell type enrichments in each spot. Firstly, we curated a set of gene signatures of common cell types in liver cancer based on the Xcell signatures (*35*) and biology prior knowledge (Table S3). Then, we defined the average log-transformed normalization expression values of the genes in the signature as the corresponding cell type scores. Taking advantage of these scores, the cell type relative enrichment degree across different tissue regions can be compared. By testing on some single cell RNA-seq datasets of liver cancer, we proved that our curated gene signatures had high sensitivity and specificity.

Furthermore, we also verified the performance of our method by comparing with the multimodal intersection analysis (MIA) (*16*), which determined the cell type enrichment degrees by performing hypergeometric test on the overlap between the tissue region-specific genes of ST data and the cell type-specific genes of single cell data. Here, we took advantage of cell type annotation and differential expression gene results of a liver cancer single cell dataset (*19*) and performed MIA on the clusters of our ST data, so that we can use the p-values of hypergeometric test to measure the enrichment of different cell types in each cluster (fig. S2C). By comparing these enrichment degrees and the mean values of our signature-based cell type scores of the all ST clusters, we observed generally high correlation (fig. S2D), which proved the reliability of our signature-based cell type scoring method. At the same time, it had the advantage of not requiring single cell data, which was more flexible.

### Intratumor spatial heterogeneity measurement

To measure the degree of intratumor heterogeneity from two aspects of transcriptome and tissue space, we proposed two metrics, transcriptome diversity degree and spatial continuity degree.

For the transcriptome diversity degree, we firstly calculated the Pearson correlation coefficients between each pair of tumor region spots based on the highly variable genes. Then we defined the sample’s transcriptome diversity degree as the 1.4826 times median absolute deviation (MAD) of these correlations, which was an approximation of standard deviation, but can avoid the impact of outliers. The larger this metric meant that the similarities among the sample’s tumors spots had larger variance, so that the sample had higher intratumor heterogeneity. Formulaically, it can be calculated as

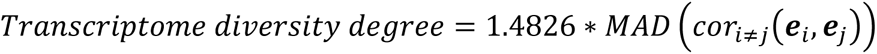

where *e_i_* indicated the expression vector of the tumor region spot *i*, and the MAD was defined as

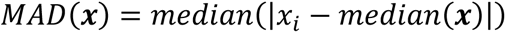

For the spatial continuity degree, we first compared the cluster identities of each tumor region spot with its six neighbor spots. Then the total fraction of the neighbor spots with the same cluster identity was defined as the spatial continuity degree. This metric measured the tumor region’s spatial heterogeneity. The larger this metric meant the sample’s tumor region more tended to be block-like (higher spatial continuity degree and lower spatial mixed degree). Formulaically, it can be calculated as

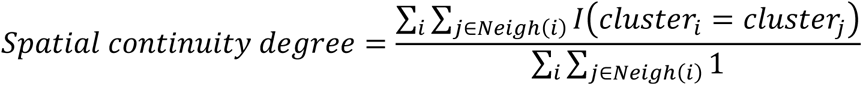

where *i* indicated a tumor region spot, and *I()* was the indicative function.

### Gene set variation analysis (GSVA)

The pathway activities of tumor cluster spots were quantified by applying gene set variation analysis (GSVA), implemented in the GSVA package (*36*). In detail, the log-transformed normalization expression matrix of tumor spots was inputted into the “gsva” function with the default parameters setting. The set of 50 cancer hallmark signatures (MSigDB, H sets), 189 oncogenic signatures (MSigDB, C6 sets) (*59*), and 96 metabolic pathways (*37*) were used to analyze. Besides, to compare the tumor clusters across patients at pathway level, we averaged the resulting GSVA score matrixes over each cluster and performed hierarchical clustering on them with Ward’s minimum variance method (Fig. 4A).

### Spatial gradient change analysis

The spatial gradient distributions of hallmark pathway activities were analyzed on our leading-edge samples (L-sections) and the intact HCC nodule (HCC-5).

For the leading-edge samples, we focused on analyzing the gradient changes from capsules or tumor-normal boundary lines to the both tumor and normal sides. The capsule (HCC-1L, 3L, 4L) boundaries were determined based on the clustering results and fine-adjusted manually by using the software Loupe Browser. When the capsules didn’t exist (ICC-1L) or were incomplete (HCC-2L, cHC-1L), the tumor-normal boundaries were decided manually according to the interface of clusters and the H&E staining images in Loupe Browser. Then, we divided the normal and tumor regions into continuous zones parallel to the shape of the boundary lines at intervals of 5 spots (fig. S3A). And the gradient changes along these zones were analyzed.

### Tumor cluster malignancy comparison analysis

To evaluate the relative malignancy degree of different ST tumor clusters, we used two liver cancer bulk datasets (TCGA-LIHC and LCI cohorts) (*39*) for comparison, which were downloaded from the HCCDB website (i.e. HCCDB15 and HCCDB6 datasets) (*38*). To reduce the impact of disease-irrelevant deaths, we truncated the patients’ survival times to five years and set their statuses as “alive” when they had longer survival times. Then, for each sample’s ST tumor cluster, we excluded its outliers and represented them by their average expression values of the remaining spots (the details were the same as the method “Cluster similarity analysis”). This step can be regarded as transforming each tumor cluster into a pseudo-bulk sample. Next, to compare the malignancy of any pair of two ST tumor clusters, we calculated the Spearman correlation coefficients between each cluster and the bulk samples (TCGA-LIHC or LCI cohorts) across a list of about 1,000 survival-related genes from HCCDB. According to the correlations, we determined which ST tumor cluster the bulk samples were more similar to, so that these bulk samples can be classified into two groups. By plotting Kaplan-Meier survival curves and performing log-rank test on these two groups, we decided the relative malignancy degrees between these two ST tumor clusters (Fig. 4C and fig. S4).

### Cluster interaction analysis

We used HCC-1T to explore the interaction between two neighbor tumor clusters, because the three tumor clusters (2, 5, and 6) in HCC-1T had clear interfaces between each other and were highly heterogeneous. For each pair of neighbor tumor clusters, we selected their interface regions with 4 spots wide (2 spots wide for each cluster) and excluded the spots identified as stromal clusters (Fig. 4D). Then, we used the CellPhoneDB (*40, 41*) to analyze the interaction strengths, which were defined as the means of the average expression level of ligand and receptor in the corresponding cluster interface spots. For each ligand-receptor pair in each interaction analysis, we performed 1,000 randomized permutations for spots’ cluster labels and recalculated the mean values, which can be seen as a null distribution. By calculating the proportion of these mean values which exceed the actual interaction strength, we obtained a p-value to measure the statistical significance of the interaction on the interface of two tumor clusters (Fig. 4E).

### Copy number variation (CNV) comparison analysis

The CNVs of each tumor spot were estimated based on their transcriptome profiles by using the method of infercnv (*50*). Firstly, for each patient, we defined the normal hepatocyte spots in their N-section as normal references. Then, all the analyzed genes were sorted by their location in the chromosomes and a sliding window of 100 genes were applied on them to calculate their moving average expression values, so that the initial copy numbers were estimated. By subtracting the normal reference copy number profiles from that of the tumor cluster spots, we got the CNV estimation of tumor spots. To reduce the impact of dropout, we took advantage of the spots’ shared-nearest-neighbor (SNN) relationships and smoothed each spot’s CNVs further by calculating the weighted average of it and its SNNs (*33*).

To confirm the CNV results inferred from ST data, we also performed the bulk WES on the PBMCs, normal sections, tumor sections, and normal/tumor regions of the leading-edge sections of the corresponding patients. Then, the copy number variations of the tumor bulk samples were called from the paired tumor-normal WES data using CNVkit software (stable version) (*51*). The normal reference adopting the PBMC or the normal section data can generate similar results. Besides, for the derived log2 copy-ratio results, the outliers were detected and filtered.

## Acknowledgements

The authors acknowledge the members of the International Co-operation Laboratory on Signal Transduction and Genergy Biotechnology (Shanghai) Co., Ltd. for excellent technical assistance. Thanks to the operating room of Eastern Hepatobiliary Surgery Hospital for providing human tumor specimens.

## Author contributions

R. Wu, W. Guo, X. Qiu contributed equally and developed the concept and discussed experiments; R. Wu and X. Qiu performed all experiments, and wrote the manuscript; W. Guo performed ST data analysis and wrote the manuscript; R. Wang performed WES data analysis; Dr. C Sui, Z. Yang provided human specimens and clinical information; Q. Lian, S, J Wu, Y Shan, S Yang, T. Wu, K. Wang, Y. Zhu, S. Wang, C Liu, Q. Zhang-Yang, B. Zheng, Z. Li, Y. Zhang, S. Shen, Y. Zhao, W. Wang, J. Bao, J. Hu, X. Wu, X Jiang provided technical assistance; Dr. H. Wang, Dr. J. Gu, and Dr. L. Chen designed research, supervised the study, guided the discussion and revised the manuscript.

## Data availability

The accession number for the raw sequencing data reported in this paper is Genome Sequence Archieve (GSA): HRA000437. Details of the ST and WES data have been deposited in the https://bigd.big.ac.cn/gsa-human/browse/HRA000437 (temporary link for review only, https://bigd.big.ac.cn/gsa-human/s/s964BDz7).

---

**Figure. S1.**
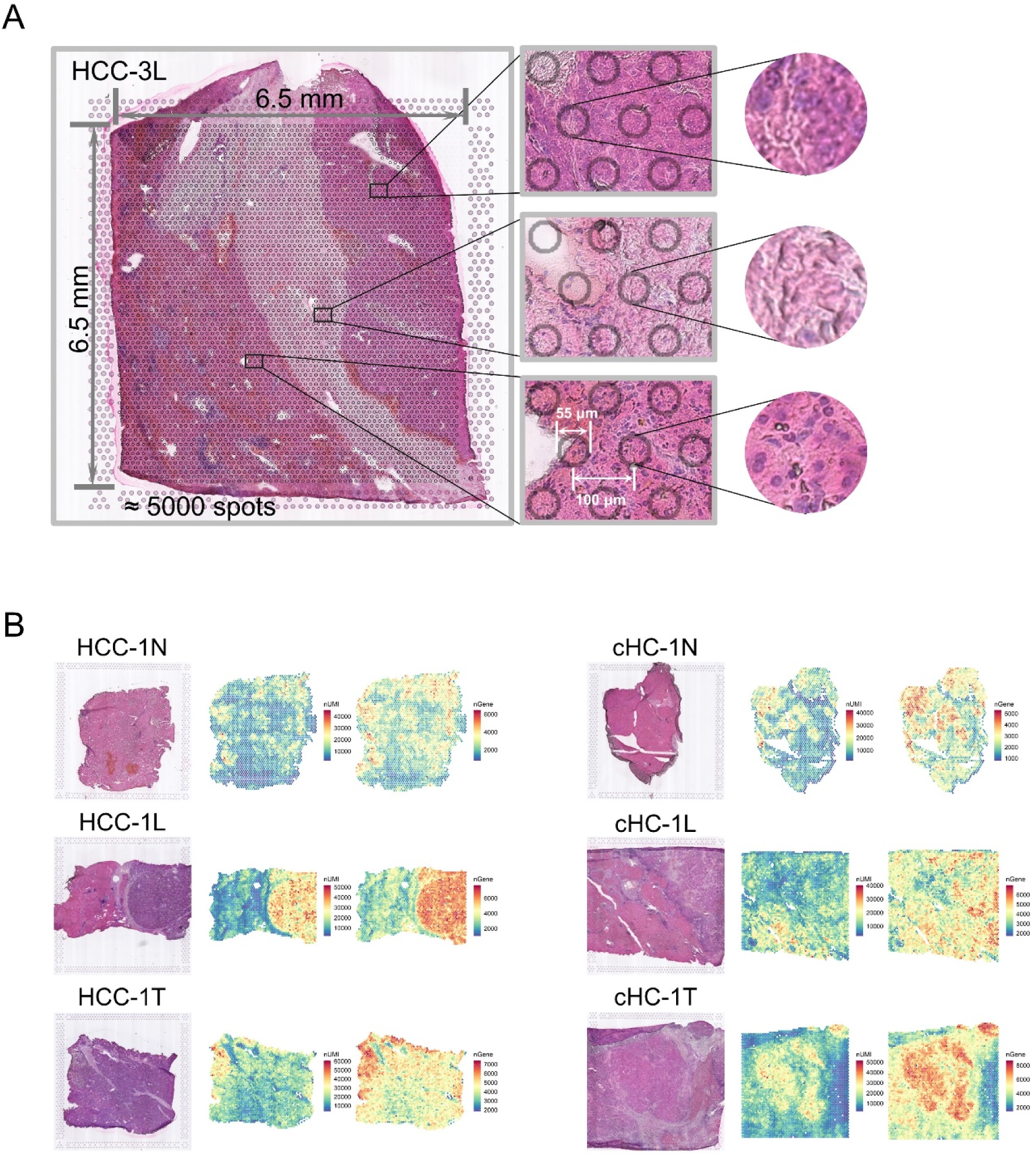
Sampling process and ST sequencing. (**A**) Spatial transcriptomics technology can detect ∼5,000 spatially barcoded spots of 55 μm diameter and 100 μm center-to-center distance in a capture area (6.5*6.5 mm^2^). (**B**) Spatial feature plots of the number of expressed transcripts (nUMIs) and genes (nGene).

**Figure. S2.**
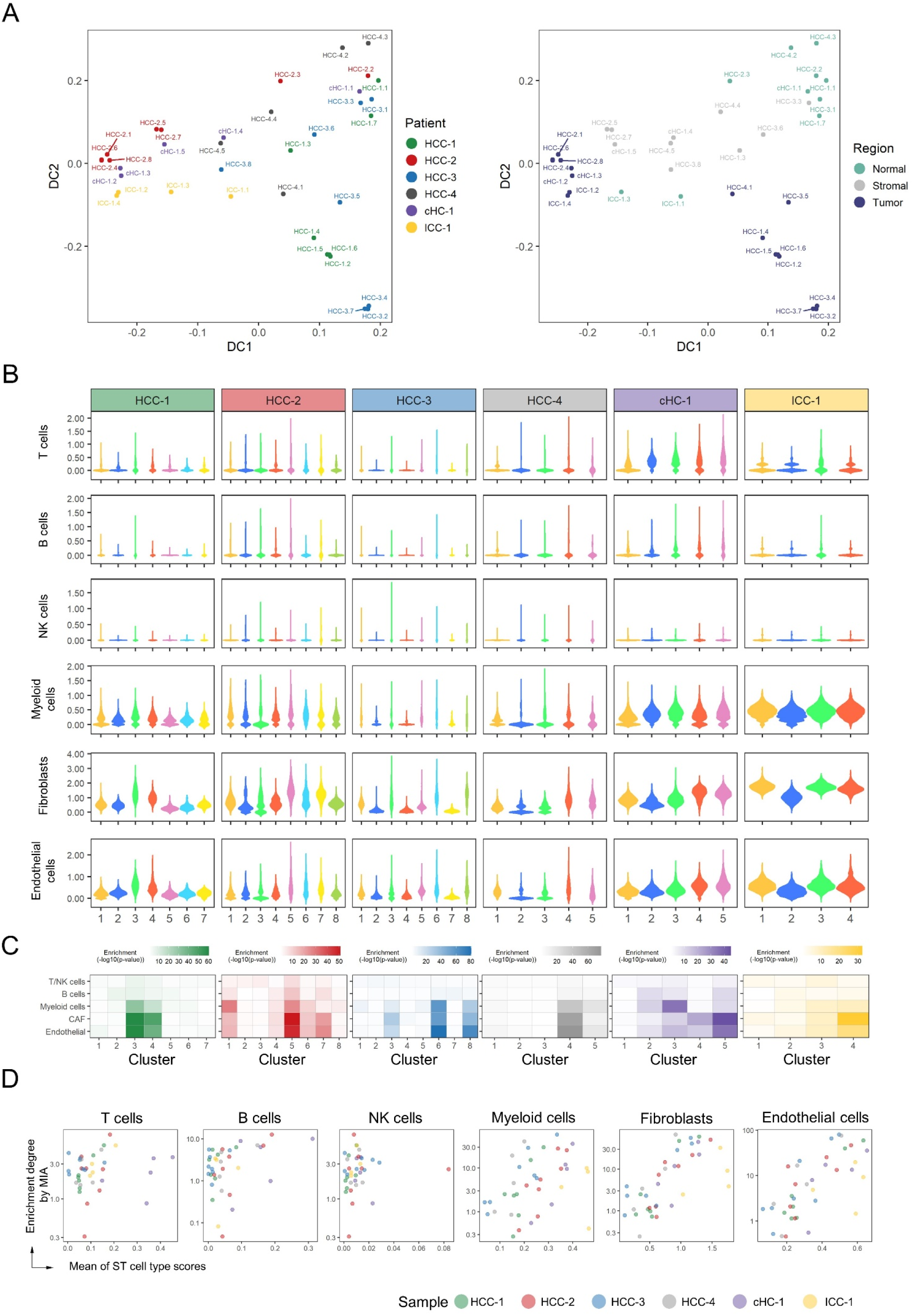
Cluster comparison and the cell type scoring. (**A**) Diffusion map of the clusters across different patient, showing the first two diffusion components. The clusters (dots) were colored by patient sources (left) and tissue regions (right) information. HCC-1.1 represented the cluster-1 of HCC-1. (**B**) Violin plots of the six stromal and immune cell type scores in each cluster. (**C**) The MIA maps of the ST defined clusters and the single cell identified cell types from a published HCC scRNA-seq dataset. Each element in the heatmap indicated the enrichment degree (-log10(p-value) of hypergeometric test) of cell types in the ST clusters, which were measured by testing on the overlap of their differential expression genes. (**D**) Comparison between the mean of ST signature-based cell type scores and the enrichment degrees by MIA. Each dot indicated one ST cluster.

**Figure S3.**
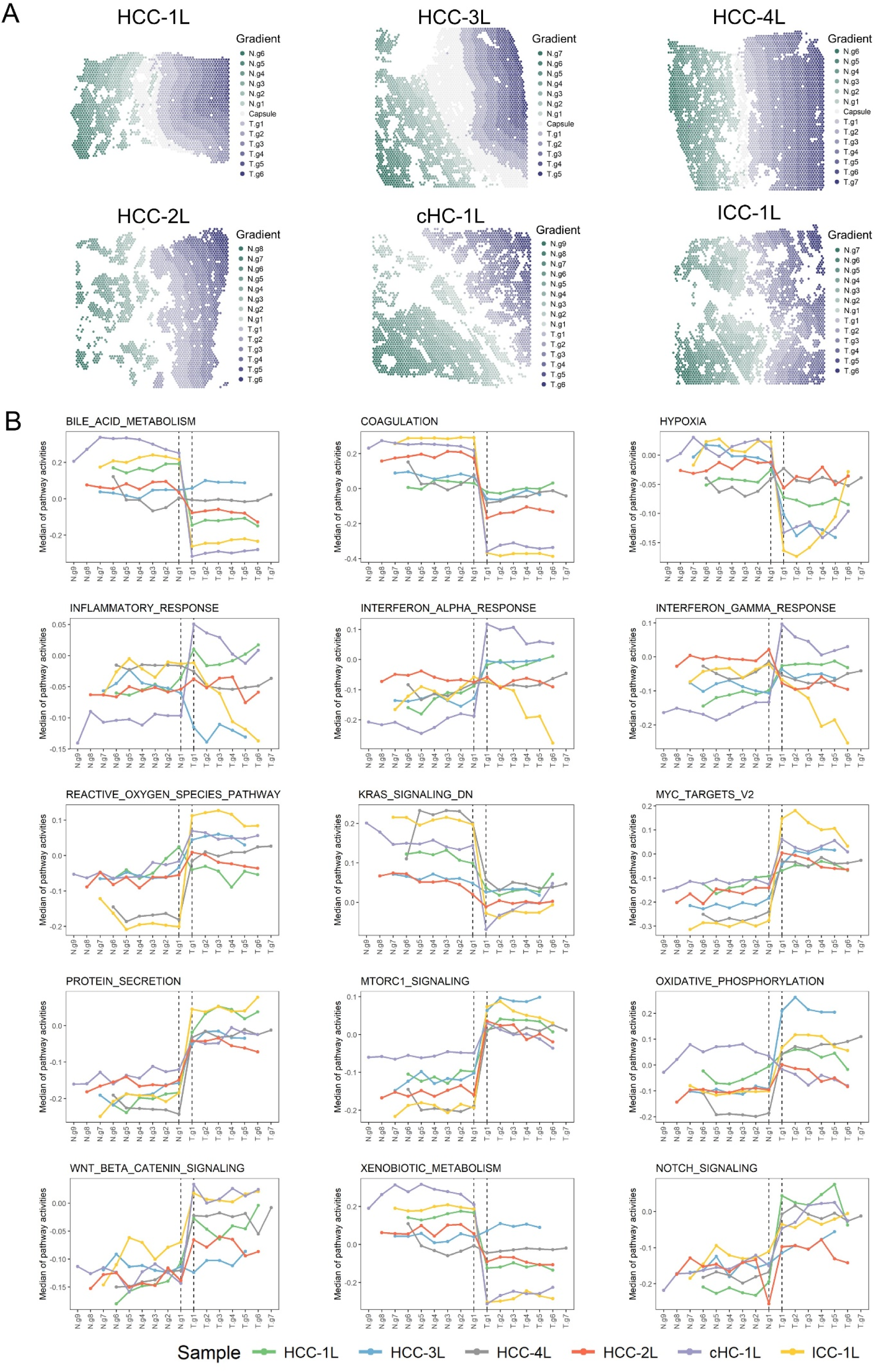
Mapping the changes of hallmark pathway activities on both sides of the transition region. (**A**) Gradient area division results on both sides of the transition region with the interval of 5 spots in L-sections. (**B**) The changes of hallmark pathways’ activities along the gradient divisions on both of the tumor and normal sides. Each dot indicated the median of the pathway activities in the corresponding area.

**Figure. S4.**
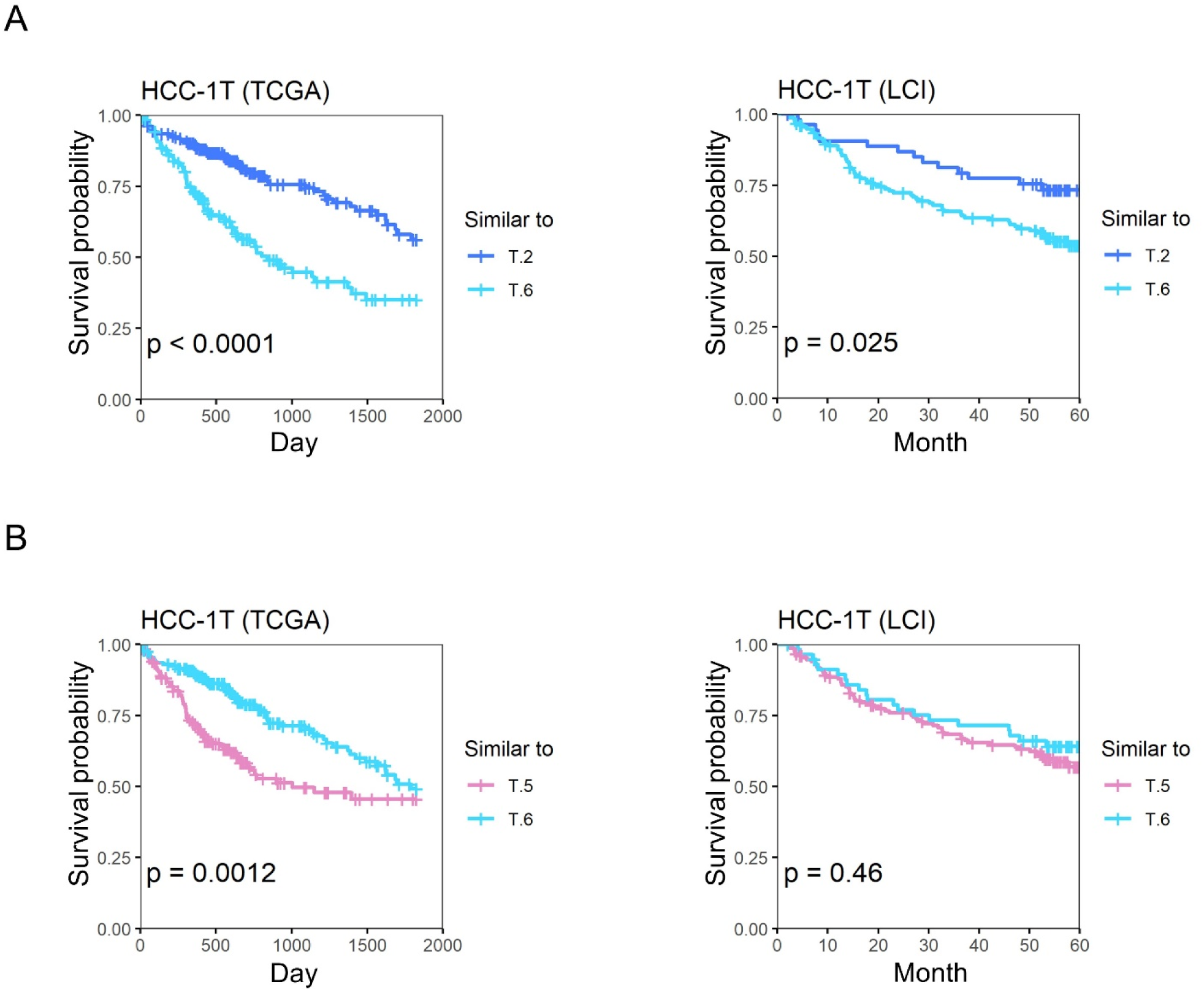
Comparison between tumor clusters. (**A**-**B**) The survival curves of two groups of patients in TCGA and LCI cohorts to compare the relative malignancy of ST tumor cluster pairs (cluster-2 vs 6, and cluster-5 vs 6 in HCC-1T). These two groups were divided according to which ST tumor cluster the bulk samples were more similar to at expression level. Log rank test was used to measure the statistical significance of their relative malignancy degrees.

**Figure. S5.**
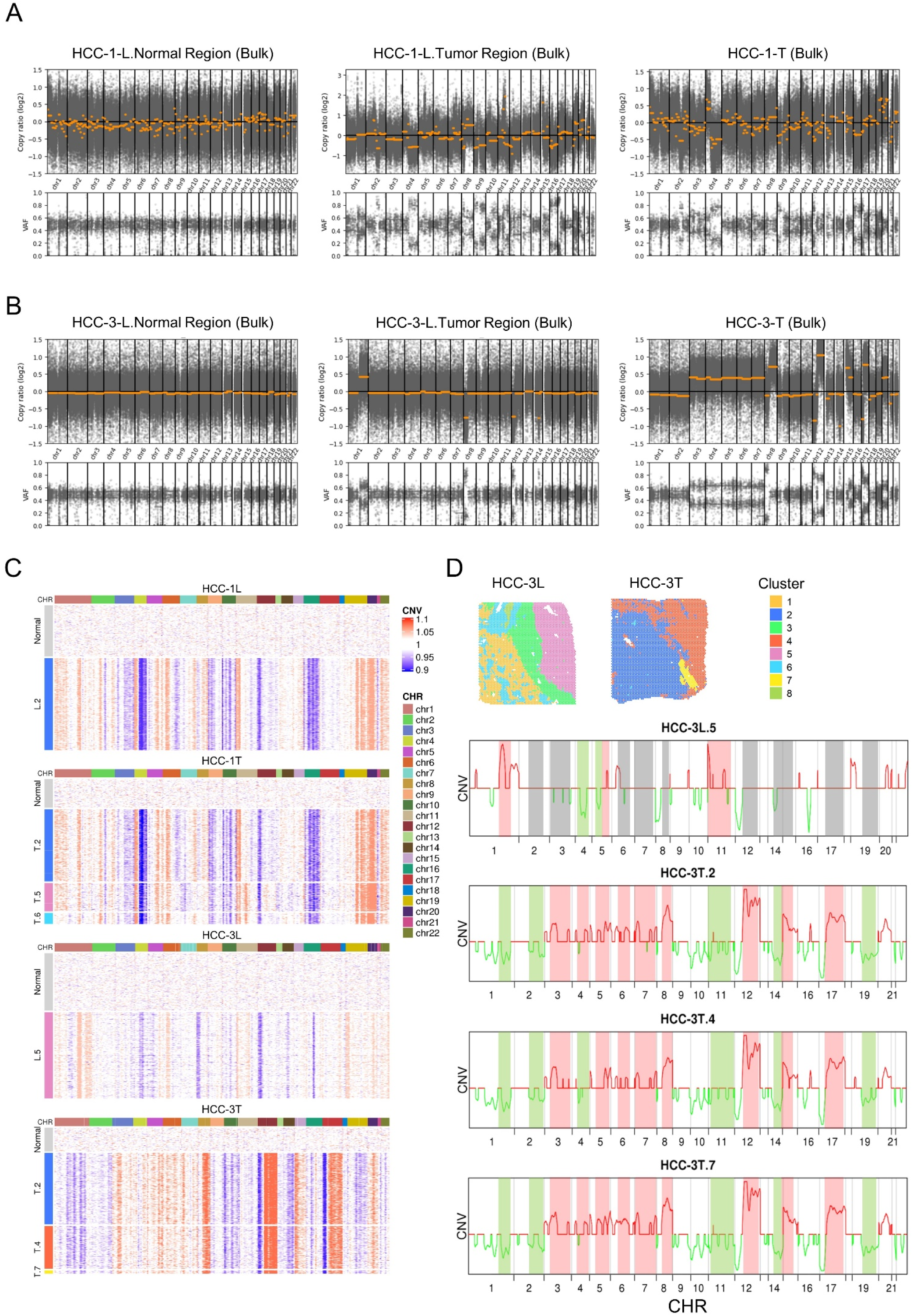
CNV Profiles of HCC-1 and HCC-3. (**A-B**) CNV profiles of the bulk samples of HCC-1 and HCC-3 patients. The three columns indicated the samples from different tissue region: the normal region of the leading-edge sections (left), the tumor region of the leading-edge sections (middle), and the tumor sections (right). (**C**) Heatmap of the inferred CNV profiles for tumor cluster spots (row). Red: amplifications; blue: deletions. The CNVs of normal references from N-sections were also presented at the top. (**D**) The averaged CNV profiles for each tumor cluster in HCC-3, inferred from spatial transcriptomes.

**Table S1.** Clinical and Pathological Data of Each Patient

| Patient ID | HCC-1 | HCC-2 | HCC-3 | HCC-4 | HCC-5 | cHC-1 | ICC-1 |
| --- | --- | --- | --- | --- | --- | --- | --- |
| Sex | Male | Female | Male | Male | Female | Female | Male |
| Age | 54 | 55 | 61 | 60 | 52 | 69 | 53 |
| Etiology of liver disease | HBV | HBV | non-HBV | HBV | HBV | HBV | non-HBV |
| Cirrhosis | Yes | No | No | No | Yes | Yes | No |
| Preoperative ALT level(U/L) | 47 | 49 | 31 | 17 | 29 | 206 | 27 |
| Preoperative AST level(U/L) | 53 | 40 | 23 | 23 | 19 | 250 | 20 |
| Preoperative serum bilirubin(umol/L) | 12.7 | 11.6 | 15.9 | 10.7 | 13.7 | 25.3 | 7.4 |
| Preoperative serum ALB(g/L) | 40.7 | 46.5 | 43 | 47.6 | 48.9 | 41.1 | 41.7 |
| Hepatic encephalopathy | 0 | 0 | 0 | 0 | 0 | 0 | 0 |
| Ascites | 0 | 0 | 0 | 0 | 0 | 0 | 0 |
| Prothrombin time (s) | 12.8 | 11.8 | 11.7 | 11.1 | 12.3 | 14.4 | 12 |
| Child-Pugh grade | A | A | A | A | A | A | A |
| BCLC stage | B | C | B | A | 0 | B | B |
| Tumor size | 6.5*6 cm | 4.3*4 cm | 8*7 cm | 3.8*3.5 cm | 1.5*1 cm | 5.0*4.2 cm | 7.6*6 cm |
| Resection margin | > 1 cm | >1 cm | > 1 cm | > 1 cm | > 1 cm | > 1 cm | > 1 cm |
| MVI | M1 | M2 | M2 | M0 | M0 | M0 | M0 |
| Histology Type | Macrotrabecular, grade III | Macrotrabecular, grade III | Macrotrabecular, grade III | Macrotrabecular, grade III | Macrotrabecular, grade III | Small bile duct type | - |
| Capsule | Complete | Incomplete | Complete | Complete | No | Incomplete | No |
| Type of Resection | R0 | R0 | R0 | R0 | R0 | R0 | R0 |
| TNM Groups | T2N0M0 | T4N0M0 | T2N0M0 | T1bN0M0 | T1aN0M0 | T1bN0M0 | T1bN1M0 |
| Tumor Stage | II | IIIb | II | Ib | Ia | Ib | IIIb |
| **Abbreviations:** ALT, Alanine transaminase; AST, Aspartate aminotransferase; ALB, Albumin; BCLC stage, Barcelona clinic liver cancer stage; MVI, Microscopic vascular invasion; TNM stage, Tumor node metastasis stage |  |  |  |  |  |  |  |

**Table S2.** Sample Spatial Transcriptomics Sequencing Data Summary

| Sample | Number of Spots Under Tissue | Median Genes per Spot | Number of Reads | Valid Barcodes | Valid UMIs | Mean Reads per Spot | Reads Mapped to Genome | Fraction Reads in Spots Under Tissue | Total Genes Detected | Median UMI Counts per Spot | Fraction of Spots Under Tissue |
| --- | --- | --- | --- | --- | --- | --- | --- | --- | --- | --- | --- |
| HCC-1N | 2956 | 2651 | 3.9E+08 | 0.97 | 1.00 | 132095.87 | 0.82 | 0.91 | 20597 | 9822.5 | 0.59 |
| HCC-1L | 2791 | 4017 | 2.9E+08 | 0.97 | 1.00 | 103838.70 | 0.91 | 0.85 | 22450 | 16407 | 0.56 |
| HCC-1T | 3184 | 4790 | 3.03E+08 | 0.97 | 1.00 | 95135.53 | 0.89 | 0.81 | 22489 | 19423.5 | 0.64 |
| cHC-1N | 2207 | 2732 | 2.72E+08 | 0.96 | 0.99 | 123233.01 | 0.75 | 0.88 | 20344 | 10753 | 0.44 |
| cHC-1L | 4516 | 4193 | 4.29E+08 | 0.96 | 0.99 | 94983.82 | 0.74 | 0.97 | 23101 | 12683.5 | 0.90 |
| cHC-1T | 4779 | 5019 | 3.88E+08 | 0.96 | 1.00 | 81139.84 | 0.84 | 0.99 | 24111 | 15239 | 0.96 |
| HCC-2N | 4628 | 1405 | 2.36E+08 | 0.97 | 1.00 | 51066.87 | 0.89 | 0.95 | 19712 | 4476.5 | 0.93 |
| HCC-2L | 4672 | 2876 | 3.18E+08 | 0.95 | 0.99 | 68158.69 | 0.74 | 0.96 | 22130 | 6604 | 0.94 |
| HCC-2T | 4733 | 3366 | 1.86E+08 | 0.96 | 0.99 | 39215.32 | 0.85 | 0.97 | 22534 | 8433 | 0.95 |
| HCC-2P | 4666 | 3059 | 1.28E+08 | 0.95 | 0.99 | 27443.02 | 0.80 | 0.97 | 21898 | 6774.5 | 0.93 |
| HCC-3N | 4289 | 2905 | 4.3E+08 | 0.96 | 0.99 | 100330.50 | 0.82 | 0.96 | 21513 | 11340 | 0.86 |
| HCC-3L | 4758 | 3043.5 | 4.32E+08 | 0.96 | 0.99 | 90739.48 | 0.85 | 0.98 | 22487 | 9348.5 | 0.95 |
| HCC-3T | 4456 | 4058 | 3.82E+08 | 0.96 | 1.00 | 85674.06 | 0.82 | 0.98 | 21259 | 20238.5 | 0.89 |
| HCC-4N | 4397 | 622 | 80540768 | 0.97 | 1.00 | 18317.21 | 0.71 | 0.94 | 19123 | 1223 | 0.88 |
| HCC-4L | 4113 | 3861 | 3.39E+08 | 0.96 | 0.99 | 82302.94 | 0.86 | 0.94 | 21928 | 12919 | 0.82 |
| HCC-4T | 4162 | 4276.5 | 3.05E+08 | 0.96 | 0.99 | 73264.38 | 0.87 | 0.94 | 21815 | 16382 | 0.83 |
| ICC-1L | 4654 | 4647.5 | 3.83E+08 | 0.96 | 0.99 | 82342.43 | 0.83 | 0.97 | 23303 | 14791 | 0.93 |
| HCC-5A | 3460 | 2024.5 | 1.2E+08 | 0.95 | 1.00 | 34576.08 | 0.84 | 0.88 | 20137 | 4954.5 | 0.69 |
| HCC-5B | 3958 | 1406 | 1.06E+08 | 0.95 | 1.00 | 26796.10 | 0.79 | 0.89 | 20075 | 2775 | 0.79 |
| HCC-5C | 3777 | 1501 | 92917728 | 0.96 | 1.00 | 24600.93 | 0.74 | 0.84 | 19984 | 3651 | 0.76 |
| HCC-5D | 4352 | 1473.5 | 1.09E+08 | 0.96 | 1.00 | 24956.50 | 0.89 | 0.96 | 20282 | 3716 | 0.87 |

**Abbreviations:** UMI, Unique Molecular Identifier.

**Table S3.** Signature Genes of Indicated Cell Types

| Cell type | Genes |
| --- | --- |
| T cells | *CD3D, CD3E, CD3G* |
| B cells | *CD19, MS4A1, CD79A* |
| NK cells | *NCAM1, KLRF1, NCR1, KLRC1* |
| Myeloid cells | *ITGAX, CD33, CEACAM8, CD68, CD163* |
| Fibroblasts | *COL1A2, FAP, PDPN, DCN, COL3A1, COL6A1, COL1A1* |
| Endothelial cells | *CLDN5, PECAM1, CD34, FLT1, VWF, ENG, CDH5* |
| CD8+ T-cells | *CD8A, CD8B* |
| CD8+ Tcm | *CD8A, CD8B, CTSW, GZMK, IL2RB, SH2D1A, GPR171, CRTAM, USP36, TMEM30B* |
| CD8+ Tem | *DHX8, GZMH, GZMK, LAG3, ZAP70, COLQ, RGS9, CXCR6, PVRIG, PYHIN1* |
| CD4+ T-cells | *CD2, CD3D, CD3E, CD3G, CD5, CD7, CD27, CD40LG, CTLA4, GOLGA4, TRAF1, CD96, ICOS, GPR171, TRAT1, FOXP3, SIRPG, RIC3* |
| Tregs | *CCR4, CCR8, CTLA4, GPR25, IL2RA, KCNA2, LAIR2, HS3ST3B1, MCF2L2, FOXP3* |
| CD4+ naive T-cells | *CD27, CLC, CTSW, DNAJB1, RBL2, HAUS3, ANKRD55, ZNF394, CHMP7* |
| CD4+ memory T-cells | *CD28, CD40LG, CCR4, CTLA4, GPR15, GZMA, GZMK, HMGB2, PDCD1, RBL2, CD226, GPR171, TRAT1, UBASH3A, ARHGAP15* |
| CD4+ Tcm | *CD40LG, CCR4, CCR8, CTLA4, DAB1, ERN1, GPR15, DNAJB1, KRT1, POU6F1, TPO, CDC14A, TRADD, DLEC1, ICOS, TRAT1, FXYD7, FBXL8, SIRPG, SLC4A5, ANKRD55, OBSCN* |
| CD4+ Tem | *CD40LG, CCR6, KLRB1, LTK, MCF2L2, GALNT8, TRAT1* |
| cDC | *CD1A, CD1B, CD1C, CD1E, CD80, CD86, FCER1A, GFRA2, CCL13, CCL17, CLEC10A, CD209, KCNK13* |
| iDC | *ALOX15, F13A1, CCL13, CCL17, CCL18, CCL23, CCL24, CD209* |
| aDC | *C1QA, C1QB, CD80, FPR3, HLA-DQA1, IL12B, CCL13, CCL17, CCL19, CCL22* |
| Monocytes | *AIF1, CYBB, FCAR, FCN1, MNDA, CFP, S100A12, CD163, CLEC5A, TREM1, RETN, FOLR2, MS4A6A, LILRA5* |
| Macrophages | *CHIT1, CYP19A1, HK3, MSR1, VSIG4, CLEC5A, ADAMDEC1, DNASE2B* |
| Macrophages M1 | *ITGAM, CD68, FCGR3A, FCGR2A, FCGR1A, CD80, CD86, HLA-DRB5, TLR4, TNF, IL1A, IL1B, IL12, IL12A, IL12B, IL6, NOS2* |
| Macrophages M2 | *ITGAM, CD68, CD163, MRC1, IL10, TGFB1, IDO1, IL8, CCL2* |
| Neutrophils | *CLC, CSF3R, FCGR3B, FPR2, HBB, CXCR2, S100A12, P2RY13, TREM1* |
| MSC | *HTR7, MMP17, PLA2G5, CDKL5, PKD2L1, ADAMTS12, ZNRF4, CTRB2* |
| ly Endothelial cells | *FLT4, HYAL2, CLEC1A, SOX18, ROBO4, CXorf36, MYCT1, KANK3* |
| mv Endothelial cells | *ACVRL1, CETP, RANGAP1, SELE, TIE1, CLDN5, VWF, HYAL2, EIF2B2, TNFSF18, LYVE1, ARHGEF15, CLEC1A, ROBO4, CXorf36, MMRN2, KANK3* |
| **Abbreviations:** Tcm, Central memory T cell; Tem, Effective memory T Cell, Treg, Regulatory T cell; cDC, conventional dendritic cell; iDC, interdigitating dendritic cell; aDC, active dendritic cell; MSC, Mesenchymal stem cell; ly Endothelial cells, lymphatic endothelial cell; mv Endothelial cells, Microvascular endothelial cell. |  |

**Table S4.**
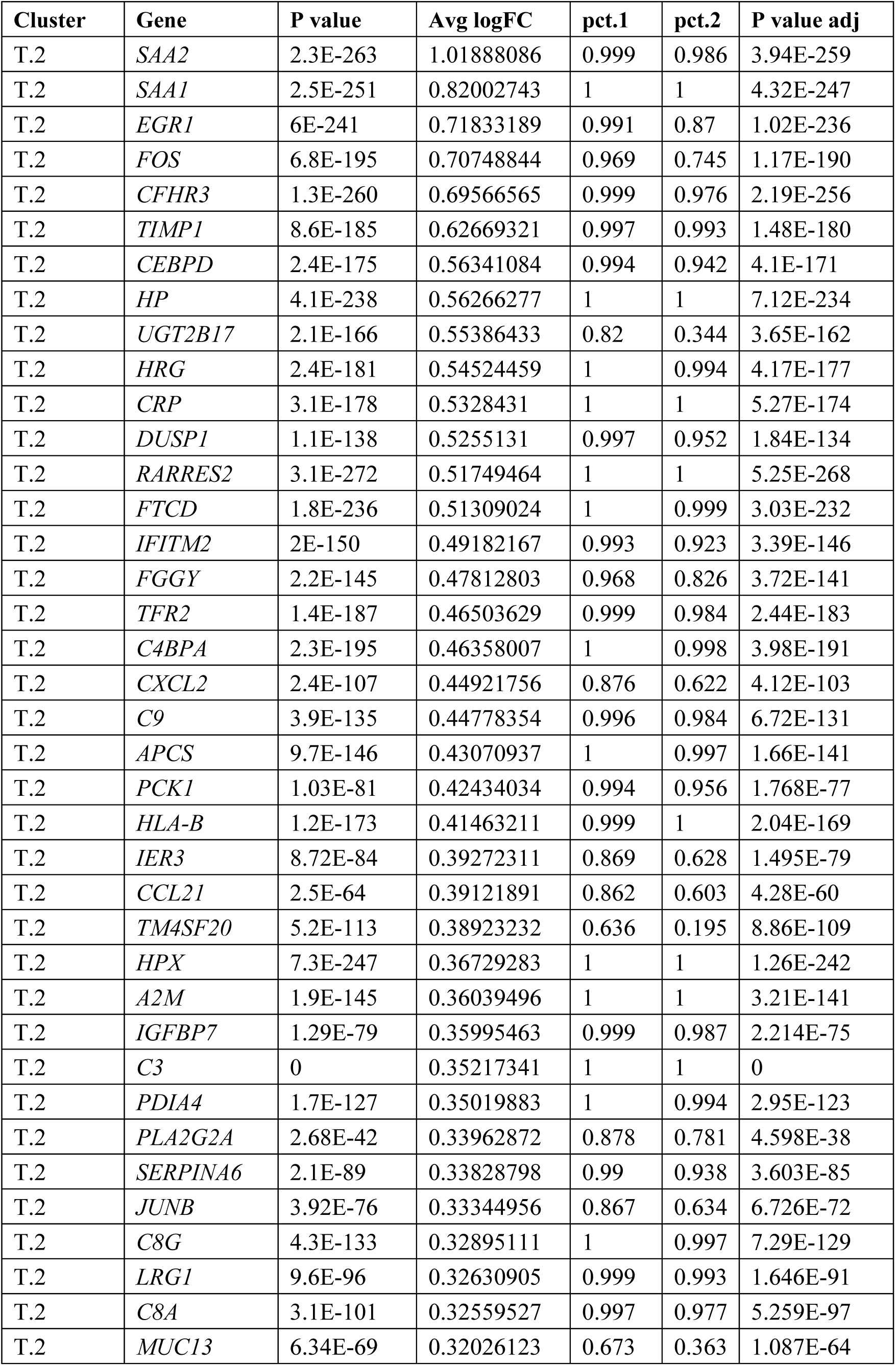
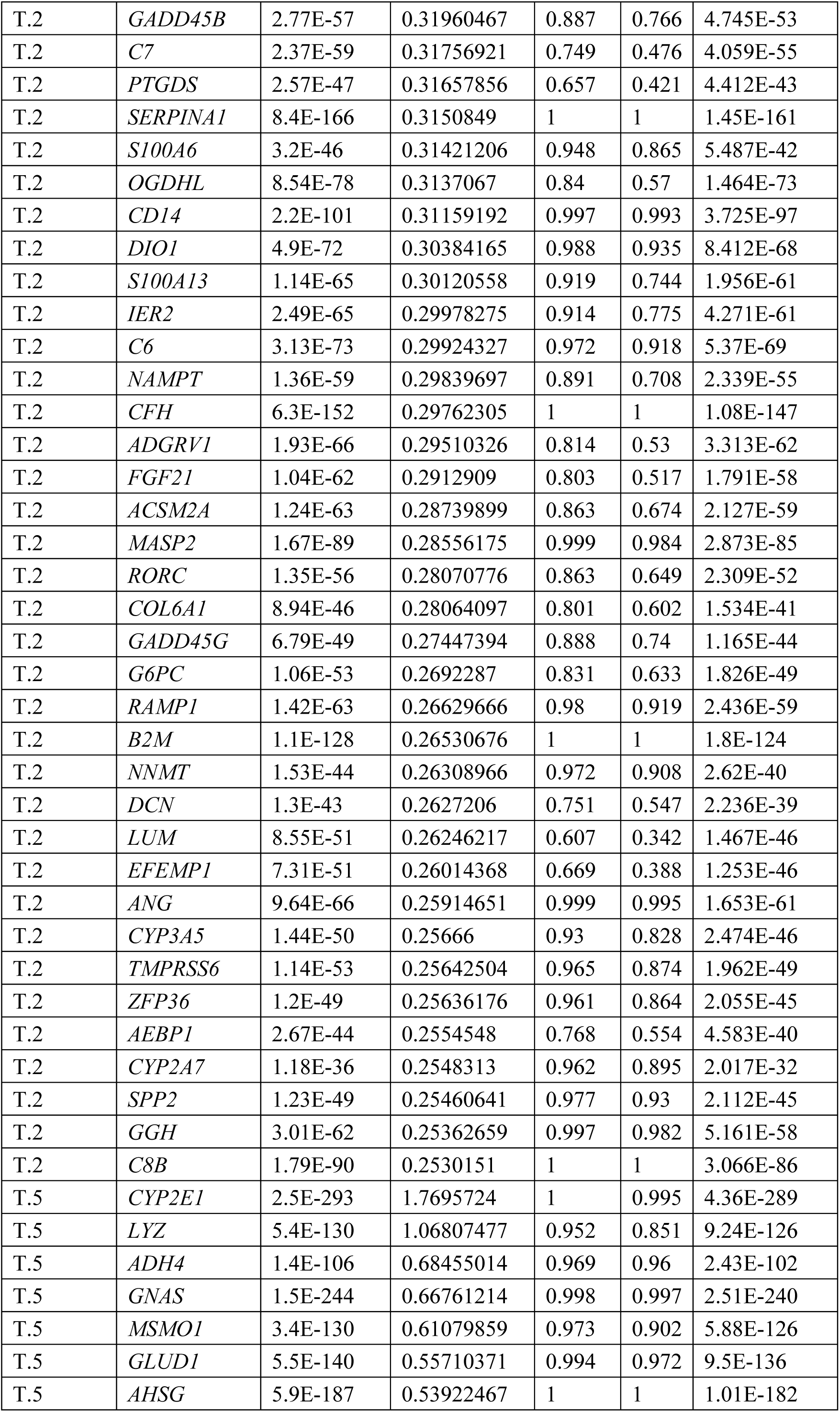
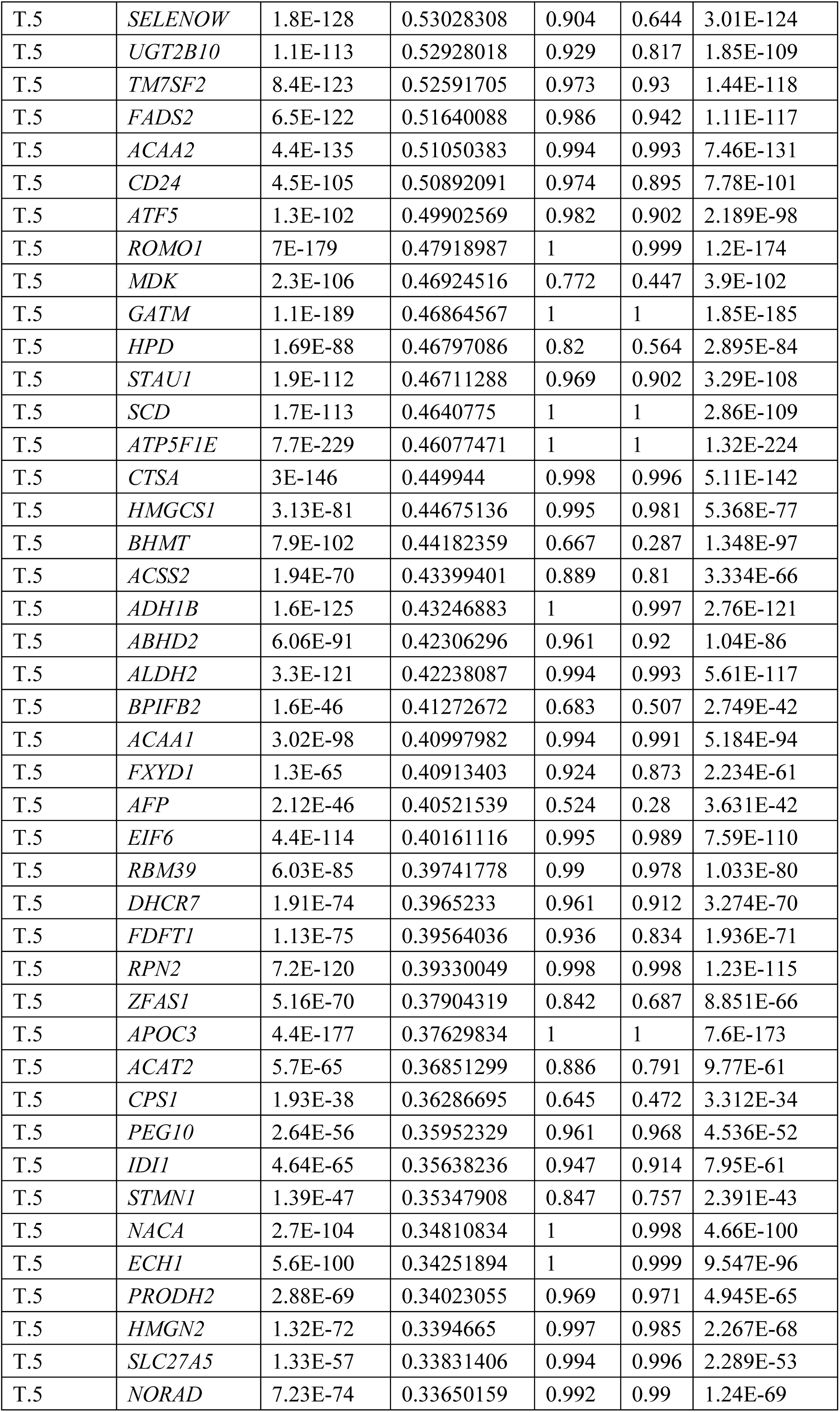
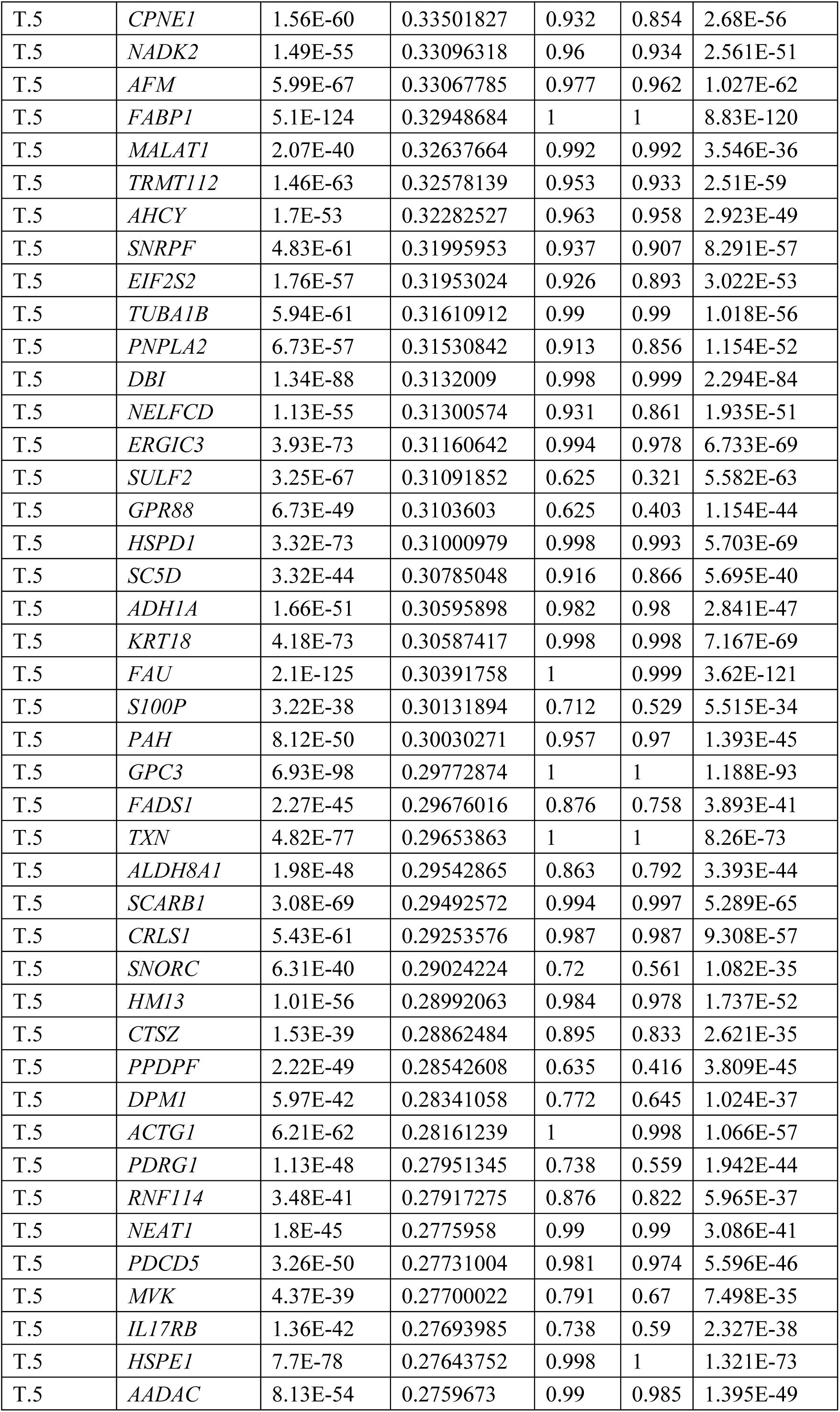
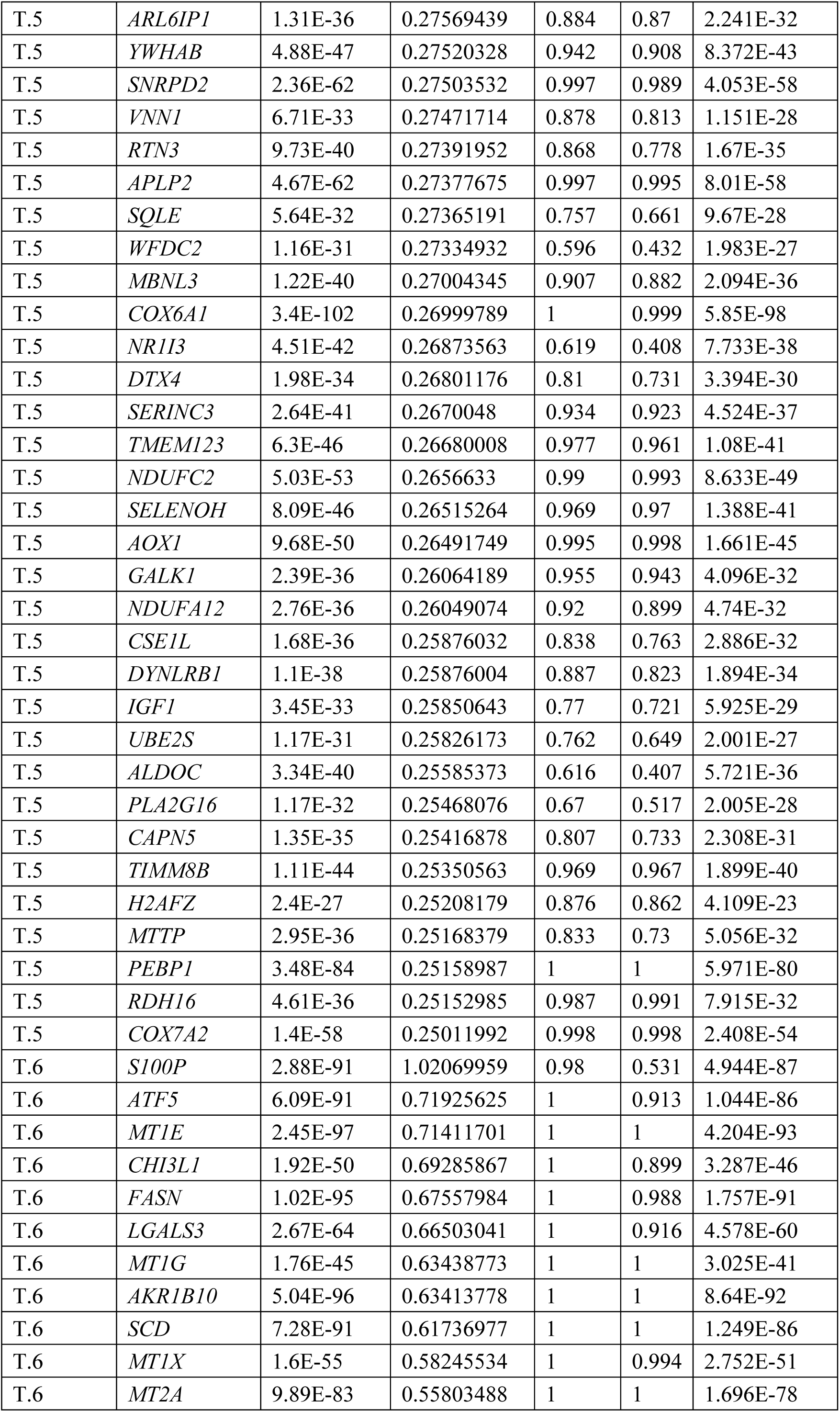
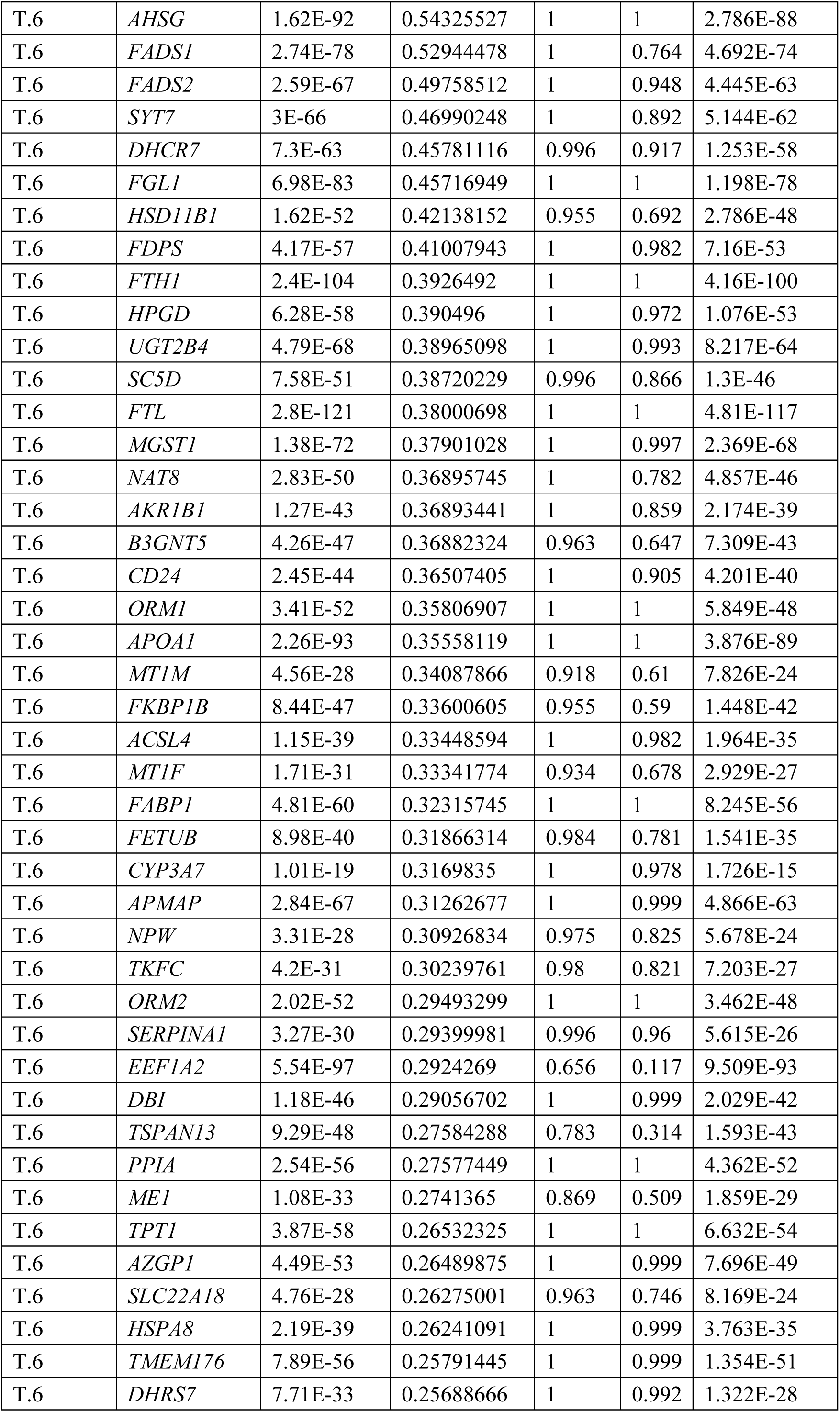
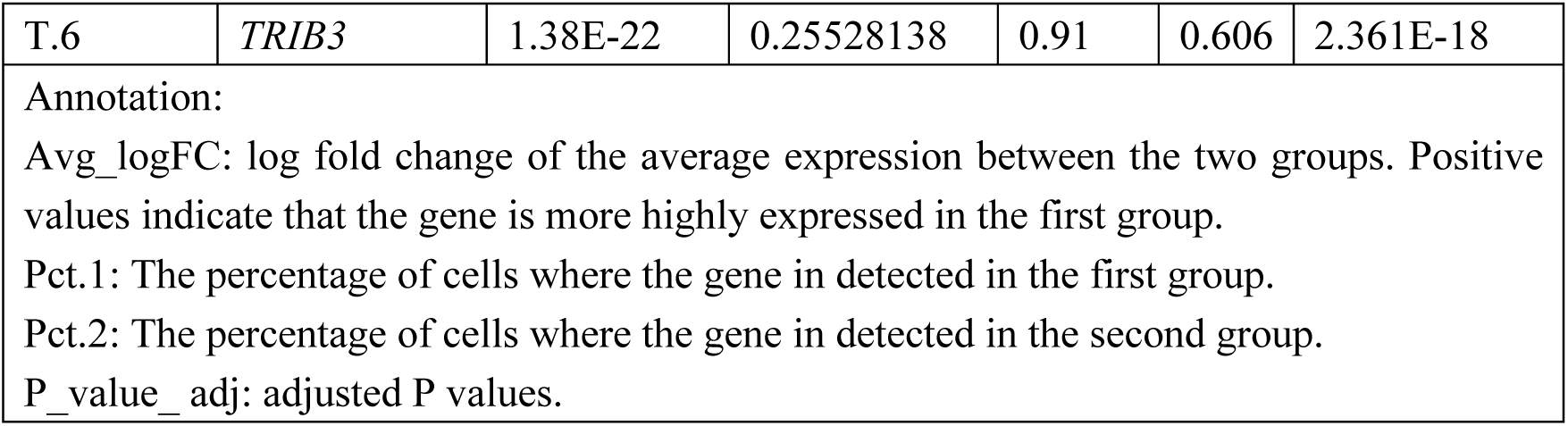
Differentially Expressed Genes of cluster-2, 5 and 6 of HCC-1T

## REFERENCES

1. I. Dagogo-Jack, A. T. Shaw, Tumour heterogeneity and resistance to cancer therapies. Nature reviews. Clinical oncology 15, 81–94 (2018).

2. N. McGranahan, C. Swanton, Clonal Heterogeneity and Tumor Evolution: Past, Present, and the Future. Cell 168, 613–628 (2017).

3. D. A. Lawson, K. Kessenbrock, R. T. Davis, N. Pervolarakis, Z. Werb, Tumour heterogeneity and metastasis at single-cell resolution. Nature Cell Biology 20, 1349–1360 (2018).

4. C. Zheng, L. Zheng, J. K. Yoo, H. Guo, Y. Zhang, X. Guo, B. Kang, R. Hu, J. Y. Huang, Q. Zhang, Z. Liu, M. Dong, X. Hu, W. Ouyang, J. Peng, Z. Zhang, Landscape of Infiltrating T Cells in Liver Cancer Revealed by Single-Cell Sequencing. Cell 169, 1342–1356.e1316 (2017).

5. N. Aizarani, A. Saviano, Sagar, L. Mailly, S. Durand, J. S. Herman, P. Pessaux, T. F. Baumert, D. Grun, A human liver cell atlas reveals heterogeneity and epithelial progenitors. Nature 572, 199–204 (2019).

6. S. Su, J. Chen, H. Yao, J. Liu, S. Yu, L. Lao, M. Wang, M. Luo, Y. Xing, F. Chen, D. Huang, J. Zhao, L. Yang, D. Liao, F. Su, M. Li, Q. Liu, E. Song, CD10(+)GPR77(+) Cancer-Associated Fibroblasts Promote Cancer Formation and Chemoresistance by Sustaining Cancer Stemness. Cell 172, 841–856 e816 (2018).

7. Z. Yin, C. Dong, K. Jiang, Z. Xu, R. Li, K. Guo, S. Shao, L. Wang, Heterogeneity of cancer-associated fibroblasts and roles in the progression, prognosis, and therapy of hepatocellular carcinoma. J Hematol Oncol 12, 101 (2019).

8. M. Zhang, H. Yang, L. Wan, Z. Wang, H. Wang, C. Ge, Y. Liu, Y. Hao, D. Zhang, G. Shi, Y. Gong, Y. Ni, C. Wang, Y. Zhang, J. Xi, S. Wang, L. Shi, L. Zhang, W. Yue, X. Pei, B. Liu, X. Yan, Single-cell transcriptomic architecture and intercellular crosstalk of human intrahepatic cholangiocarcinoma. J Hepatol 73, 1118–1130 (2020).

9. K. H. Chen, A. N. Boettiger, J. R. Moffitt, S. Wang, X. Zhuang, RNA imaging. Spatially resolved, highly multiplexed RNA profiling in single cells. Science 348, aaa6090 (2015).

10. C. L. Eng, M. Lawson, Q. Zhu, R. Dries, N. Koulena, Y. Takei, J. Yun, C. Cronin, C. Karp, G. C. Yuan, L. Cai, Transcriptome-scale super-resolved imaging in tissues by RNA seqFISH. Nature 568, 235–239 (2019).

11. D. J. Burgess, Spatial transcriptomics coming of age. Nature Reviews Genetics 20, 317–317 (2019).

12. S. Vickovic, G. Eraslan, F. Salmén, J. Klughammer, L. Stenbeck, D. Schapiro, T. ij, R. Bonneau, L. Bergenstr?hle, J. F. Navarro, J. Gould, G. K. Griffin, Borg, M. Ronaghi, J. Frisén, J. Lundeberg, A. Regev, P. L. St?hl, High-definition spatial transcriptomics for in situ tissue profiling. Nature methods 16, 987–990 (2019).

13. P. L. Stahl, F. Salmen, S. Vickovic, A. Lundmark, J. F. Navarro, J. Magnusson, S. Giacomello, M. Asp, J. O. Westholm, M. Huss, A. Mollbrink, S. Linnarsson, S. Codeluppi, A. Borg, F. Ponten, P. I. Costea, P. Sahlen, J. Mulder, O. Bergmann, J. Lundeberg, J. Frisen, Visualization and analysis of gene expression in tissue sections by spatial transcriptomics. Science 353, 78–82 (2016).

14. K. Thrane, H. Eriksson, J. Maaskola, J. Hansson, J. Lundeberg, Spatially Resolved Transcriptomics Enables Dissection of Genetic Heterogeneity in Stage III Cutaneous Malignant Melanoma. Cancer research 78, 5970–5979 (2018).

15. E. Berglund, J. Maaskola, N. Schultz, S. Friedrich, M. Marklund, J. Bergenstråhle, F. Tarish, A. Tanoglidi, S. Vickovic, L. Larsson, F. Salmén, C. Ogris, K. Wallenborg, J. Lagergren, P. Ståhl, E. Sonnhammer, T. Helleday, J. Lundeberg, Spatial maps of prostate cancer transcriptomes reveal an unexplored landscape of heterogeneity. Nature communications 9, 2419 (2018).

16. R. Moncada, D. Barkley, F. Wagner, M. Chiodin, J. C. Devlin, M. Baron, C. H. Hajdu, D. M. Simeone, I. Yanai, Integrating microarray-based spatial transcriptomics and single-cell RNA-seq reveals tissue architecture in pancreatic ductal adenocarcinomas. Nat Biotechnol 38, 333–342 (2020).

17. M. Asp, S. Giacomello, L. Larsson, C. Wu, D. Furth, X. Qian, E. Wardell, J. Custodio, J. Reimegard, F. Salmen, C. Osterholm, P. L. Stahl, E. Sundstrom, E. Akesson, O. Bergmann, M. Bienko, A. Mansson-Broberg, M. Nilsson, C. Sylven, J. Lundeberg, A Spatiotemporal Organ-Wide Gene Expression and Cell Atlas of the Developing Human Heart. Cell 179, 1647–1660 e1619 (2019).

18. D. Sia, A. Villanueva, S. L. Friedman, J. M. Llovet, Liver Cancer Cell of Origin, Molecular Class, and Effects on Patient Prognosis. Gastroenterology 152, 745–761 (2017).

19. B. Losic, A. J. Craig, C. Villacorta-Martin, S. N. Martins-Filho, N. Akers, X. Chen, M. E. Ahsen, J. von Felden, I. Labgaa, D. D’Avola, K. Allette, S. A. Lira, G. C. Furtado, T. Garcia-Lezana, P. Restrepo, A. Stueck, S. C. Ward, M. I. Fiel, S. P. Hiotis, G. Gunasekaran, D. Sia, E. E. Schadt, R. Sebra, M. Schwartz, J. M. Llovet, S. Thung, G. Stolovitzky, A. Villanueva, Intratumoral heterogeneity and clonal evolution in liver cancer. Nat Commun 11, 291 (2020).

20. T. F. Greten, X. W. Wang, F. Korangy, Current concepts of immune based treatments for patients with HCC: from basic science to novel treatment approaches. Gut 64, 842–848 (2015).

21. A. B. El-Khoueiry, B. Sangro, T. Yau, T. S. Crocenzi, M. Kudo, C. Hsu, T. Y. Kim, S. P. Choo, J. Trojan, T. H. R. Welling, T. Meyer, Y. K. Kang, W. Yeo, A. Chopra, J. Anderson, C. Dela Cruz, L. Lang, J. Neely, H. Tang, H. B. Dastani, I. Melero, Nivolumab in patients with advanced hepatocellular carcinoma (CheckMate 040): an open-label, non-comparative, phase 1/2 dose escalation and expansion trial. Lancet (London, England) 389, 2492–2502 (2017).

22. J. M. Llovet, S. Ricci, V. Mazzaferro, P. Hilgard, E. Gane, J. F. Blanc, A. C. de Oliveira, A. Santoro, J. L. Raoul, A. Forner, M. Schwartz, C. Porta, S. Zeuzem, L. Bolondi, T. F. Greten, P. R. Galle, J. F. Seitz, I. Borbath, H. u. D, T. Giannaris, M. Shan, M. Moscovici, D. Voliotis, J. Bruix, S. I. S. G. CollectiveName, Sorafenib in advanced hepatocellular carcinoma. The New England journal of medicine 359, 378–390 (2008).

23. A. X. Zhu, R. S. Finn, J. Edeline, S. Cattan, S. Ogasawara, D. Palmer, C. Verslype, V. Zagonel, L. Fartoux, A. Vogel, D. Sarker, G. Verset, S. L. Chan, J. Knox, B. Daniele, A. L. Webber, S. W. Ebbinghaus, J. Ma, A. B. Siegel, A. L. Cheng, M. Kudo, K.-i. CollectiveName, Pembrolizumab in patients with advanced hepatocellular carcinoma previously treated with sorafenib (KEYNOTE-224): a non-randomised, open-label phase 2 trial. The Lancet. Oncology 19, 940–952 (2018).

24. X. Ding, M. He, A. W. H. Chan, Q. X. Song, S. C. Sze, H. Chen, M. K. H. Man, K. Man, S. L. Chan, P. B. S. Lai, X. Wang, N. Wong, Genomic and Epigenomic Features of Primary and Recurrent Hepatocellular Carcinomas. Gastroenterology, (2020).

25. S. Gordon, P. R. Taylor, Monocyte and macrophage heterogeneity. Nature reviews. Immunology 5, 953–964 (2005).

26. G. Ishii, A. Ochiai, S. Neri, Phenotypic and functional heterogeneity of cancer-associated fibroblast within the tumor microenvironment. Advanced drug delivery reviews 99, 186–196 (2016).

27. M. D. Lynch, F. M. Watt, Fibroblast heterogeneity: implications for human disease. The Journal of clinical investigation 128, 26–35 (2018).

28. A. Sharma, J. J. W. Seow, C. A. Dutertre, R. Pai, C. Bleriot, A. Mishra, R. M. M. Wong, G. S. N. Singh, S. Sudhagar, S. Khalilnezhad, S. Erdal, H. M. Teo, A. Khalilnezhad, S. Chakarov, T. K. H. Lim, A. C. Y. Fui, A. K. W. Chieh, C. P. Chung, G. K. Bonney, B. K. Goh, J. K. Y. Chan, P. K. H. Chow, F. Ginhoux, R. DasGupta, Onco-fetal Reprogramming of Endothelial Cells Drives Immunosuppressive Macrophages in Hepatocellular Carcinoma. Cell 183, 377–394 e321 (2020).

29. B. Zheng, D. Wang, X. Qiu, G. Luo, T. Wu, S. Yang, Z. Li, Y. Zhu, S. Wang, R. Wu, C. Sui, Z. Gu, S. Shen, S. Jeong, X. Wu, J. Gu, H. Wang, L. Chen, Trajectory and Functional Analysis of PD-1(high) CD4(+)CD8(+) T Cells in Hepatocellular Carcinoma by Single-Cell Cytometry and Transcriptome Sequencing. Adv Sci (Weinh) 7, 2000224 (2020).

30. I. Korsunsky, N. Millard, J. Fan, K. Slowikowski, F. Zhang, K. Wei, Y. Baglaenko, M. Brenner, P. R. Loh, S. Raychaudhuri, Fast, sensitive and accurate integration of single-cell data with Harmony. Nat Methods 16, 1289–1296 (2019).

31. C. Y. Lin, J. Loven, P. B. Rahl, R. M. Paranal, C. B. Burge, J. E. Bradner, T. I. Lee, R. A. Young, Transcriptional amplification in tumor cells with elevated c-Myc. Cell 151, 56–67 (2012).

32. T. Stuart, A. Butler, P. Hoffman, C. Hafemeister, E. Papalexi, W. M. Mauck, 3rd, Y. Hao, M. Stoeckius, P. Smibert, R. Satija, Comprehensive Integration of Single-Cell Data. Cell 177, 1888–1902 e1821 (2019).

33. W. Guo, D. Wang, S. Wang, Y. Shan, C. Liu, J. Gu, scCancer: a package for automated processing of single-cell RNA-seq data in cancer. Briefings in Bioinformatics, (2020).

34. P. Angerer, L. Haghverdi, M. Buttner, F. J. Theis, C. Marr, F. Buettner, destiny: diffusion maps for large-scale single-cell data in R. Bioinformatics 32, 1241–1243 (2016).

35. D. Aran, Z. Hu, A. J. Butte, xCell: digitally portraying the tissue cellular heterogeneity landscape. Genome Biol 18, 220 (2017).

36. S. Hänzelmann, R. Castelo, J. Guinney, GSVA: gene set variation analysis for microarray and RNA-seq data. BMC Bioinformatics 14, 7 (2013).

37. E. Gaude, C. Frezza, Tissue-specific and convergent metabolic transformation of cancer correlates with metastatic potential and patient survival. Nat Commun 7, 13041 (2016).

38. Q. Lian, S. Wang, G. Zhang, D. Wang, G. Luo, J. Tang, L. Chen, J. Gu, HCCDB: A Database of Hepatocellular Carcinoma Expression Atlas. Genomics Proteomics Bioinformatics 16, 269–275 (2018).

39. S. Roessler, H. L. Jia, A. Budhu, M. Forgues, Q. H. Ye, J. S. Lee, S. S. Thorgeirsson, Z. Sun, Z. Y. Tang, L. X. Qin, X. W. Wang, A unique metastasis gene signature enables prediction of tumor relapse in early-stage hepatocellular carcinoma patients. Cancer Res 70, 10202–10212 (2010).

40. M. Efremova, M. Vento-Tormo, S. A. Teichmann, R. Vento-Tormo, CellPhoneDB: inferring cell-cell communication from combined expression of multi-subunit ligand-receptor complexes. Nat Protoc 15, 1484–1506 (2020).

41. R. Vento-Tormo, M. Efremova, R. A. Botting, M. Y. Turco, M. Vento-Tormo, K. B. Meyer, J. E. Park, E. Stephenson, K. Polanski, A. Goncalves, L. Gardner, S. Holmqvist, J. Henriksson, A. Zou, A. M. Sharkey, B. Millar, B. Innes, L. Wood, A. Wilbrey-Clark, R. P. Payne, M. A. Ivarsson, S. Lisgo, A. Filby, D. H. Rowitch, J. N. Bulmer, G. J. Wright, M. J. T. Stubbington, M. Haniffa, A. Moffett, S. A. Teichmann, Single-cell reconstruction of the early maternal-fetal interface in humans. Nature 563, 347–353 (2018).

42. M. Bry, R. Kivela, V. M. Leppanen, K. Alitalo, Vascular endothelial growth factor-B in physiology and disease. Physiol Rev 94, 779–794 (2014).

43. S. Soker, S. Takashima, H. Q. Miao, G. Neufeld, M. Klagsbrun, Neuropilin-1 is expressed by endothelial and tumor cells as an isoform-specific receptor for vascular endothelial growth factor. Cell 92, 735–745 (1998).

44. L. Seetharam, N. Gotoh, Y. Maru, G. Neufeld, S. Yamaguchi, M. Shibuya, A unique signal transduction from FLT tyrosine kinase, a receptor for vascular endothelial growth factor VEGF. Oncogene 10, 135–147 (1995).

45. B. D. Ferguson, R. Liu, C. E. Rolle, Y. H. Tan, V. Krasnoperov, R. Kanteti, M. S. Tretiakova, G. M. Cervantes, R. Hasina, R. D. Hseu, A. J. Iafrate, T. Karrison, M. K. Ferguson, A. N. Husain, L. Faoro, E. E. Vokes, P. S. Gill, R. Salgia, The EphB4 receptor tyrosine kinase promotes lung cancer growth: a potential novel therapeutic target. PLoS One 8, e67668 (2013).

46. M. Pietila, P. Sahgal, E. Peuhu, N. Z. Jantti, I. Paatero, E. Narva, H. Al-Akhrass, J. Lilja, M. Georgiadou, O. M. Andersen, A. Padzik, H. Sihto, H. Joensuu, M. Blomqvist, I. Saarinen, P. J. Bostrom, P. Taimen, J. Ivaska, SORLA regulates endosomal trafficking and oncogenic fitness of HER2. Nat Commun 10, 2340 (2019).

47. K. Ieguchi, Y. Maru, Roles of EphA1/A2 and ephrin-A1 in cancer. Cancer Sci 110, 841–848 (2019).

48. A. Kinoshita, T. Shah, M. M. Tangredi, D. K. Strickland, B. T. Hyman, The intracellular domain of the low density lipoprotein receptor-related protein modulates transactivation mediated by amyloid precursor protein and Fe65. J Biol Chem 278, 41182–41188 (2003).

49. H. Mao, P. Lockyer, L. Li, C. M. Ballantyne, C. Patterson, L. Xie, X. Pi, Endothelial LRP1 regulates metabolic responses by acting as a co-activator of PPARgamma. Nat Commun 8, 14960 (2017).

50. A. P. Patel, I. Tirosh, J. J. Trombetta, A. K. Shalek, S. M. Gillespie, H. Wakimoto, D. P. Cahill, B. V. Nahed, W. T. Curry, R. L. Martuza, D. N. Louis, O. Rozenblatt-Rosen, M. L. Suva, A. Regev, B. E. Bernstein, Single-cell RNA-seq highlights intratumoral heterogeneity in primary glioblastoma. Science 344, 1396–1401 (2014).

51. E. Talevich, A. H. Shain, T. Botton, B. C. Bastian, CNVkit: Genome-Wide Copy Number Detection and Visualization from Targeted DNA Sequencing. PLoS Comput Biol 12, e1004873 (2016).

52. M. Liu, L. Chen, T. H. Chan, J. Wang, Y. Li, Y. Li, T. T. Zeng, Y. F. Yuan, X. Y. Guan, Serum and glucocorticoid kinase 3 at 8q13.1 promotes cell proliferation and survival in hepatocellular carcinoma. Hepatology 55, 1754–1765 (2012).

53. J. Zucman-Rossi, A. Villanueva, J. C. Nault, J. M. Llovet, Genetic Landscape and Biomarkers of Hepatocellular Carcinoma. Gastroenterology 149, 1226–1239 e1224 (2015).

54. D. F. Quail, J. A. Joyce, Microenvironmental regulation of tumor progression and metastasis. Nat Med 19, 1423–1437 (2013).

55. T. Wu, Y. Dai, Tumor microenvironment and therapeutic response. Cancer Lett 387, 61–68 (2017).

56. Z. F. Lim, P. C. Ma, Emerging insights of tumor heterogeneity and drug resistance mechanisms in lung cancer targeted therapy. J Hematol Oncol 12, 134 (2019).

57. R. Rahmanzade, Redefinition of tumor capsule: Rho-dependent clustering of cancer-associated fibroblasts in favor of tensional homeostasis. Med Hypotheses 135, 109425 (2020).

58. A. J. Craig, J. von Felden, T. Garcia-Lezana, S. Sarcognato, A. Villanueva, Tumour evolution in hepatocellular carcinoma. Nat Rev Gastroenterol Hepatol 17, 139–152 (2020).

59. Y. Subbannayya, S. M. Pinto, K. Bosl, T. S. K. Prasad, R. K. Kandasamy, Dynamics of Dual Specificity Phosphatases and Their Interplay with Protein Kinases in Immune Signaling. International Journal of Molecular Sciences 20, (2019).

